# Molecular and functional alterations in the cerebral microvasculature in an optimized mouse model of sepsis-associated cognitive dysfunction

**DOI:** 10.1101/2024.05.28.596050

**Authors:** Paulo Ávila-Gómez, Yuto Shingai, Sabyasachi Dash, Catherine Liu, Keri Callegari, Heidi Meyer, Anne Khodarkovskaya, Daiki Aburakawa, Hiroki Uchida, Giuseppe Faraco, Lidia Garcia-Bonilla, Josef Anrather, Francis S Lee, Costantino Iadecola, Teresa Sanchez

## Abstract

Systemic inflammation has been implicated in the development and progression of neurodegenerative conditions such as cognitive impairment and dementia. Recent clinical studies indicate an association between sepsis, endothelial dysfunction, and cognitive decline. However, the investigations of the role and therapeutic potential of the cerebral microvasculature in systemic inflammation-induced cognitive dysfunction have been limited by the lack of standardized experimental models for evaluating the alterations in the cerebral microvasculature and cognition induced by the systemic inflammatory response. Herein, we validated a mouse model of endotoxemia that recapitulates key pathophysiology related to sepsis-induced cognitive dysfunction, including the induction of an acute systemic hyperinflammatory response, blood-brain barrier (BBB) leakage, neurovascular inflammation, and memory impairment after recovery from the systemic inflammatory response. In the acute phase, we identified novel molecular (e.g. upregulation of plasmalemma vesicle associated protein, a driver of endothelial permeability, and the pro-coagulant plasminogen activator inhibitor-1, PAI-1) and functional perturbations (i.e., albumin and small molecule BBB leakage) in the cerebral microvasculature along with neuroinflammation. Remarkably, small molecule BBB permeability, elevated levels of PAI-1, intra/perivascular fibrin/fibrinogen deposition and microglial activation persisted 1 month after recovery from sepsis. We also highlight molecular neuronal alterations of potential clinical relevance following systemic inflammation including changes in neurofilament phosphorylation and decreases in postsynaptic density protein 95 and brain-derived neurotrophic factor suggesting diffuse axonal injury, synapse degeneration and impaired neurotrophism. Our study serves as a standardized model to support future mechanistic studies of sepsis-associated cognitive dysfunction and to identify novel endothelial therapeutic targets for this devastating condition.

**SIGNIFICANCE:** The limited knowledge of how systemic inflammation contributes to cognitive decline is a major obstacle to the development of novel therapies for dementia and other neurodegenerative diseases. Clinical evidence supports a role for the cerebral microvasculature in sepsis-induced neurocognitive dysfunction, but the investigation of the underlying mechanisms has been limited by the lack of standardized experimental models. Herein, we optimized a mouse model that recapitulates important pathophysiological aspects of systemic inflammation-induced cognitive decline and identified key alterations in the cerebral microvasculature associated with cognitive dysfunction. Our study provides a reliable experimental model for mechanistic studies and therapeutic discovery of the impact of systemic inflammation on cerebral microvascular function and the development and progression of cognitive impairment.

## INTRODUCTION

Systemic inflammation has been associated with the development and progression of several neurodegenerative conditions, including cognitive impairment and dementia. Up to 50% of patients who recover from severe systemic inflammation (i.e. systemic inflammatory response syndrome, SIRS, and sepsis) develop persistent cognitive impairment and accelerated dementia (Annane et al., 2015; Beltran-Garcia et al., 2020; Bettcher et al., 2021; Chou et al., 2017; Hampshire et al., 2021; Holmes et al., 2009; Iwashyna et al., 2010; Liu et al., 2021; Manabe et al., 2022; Mostel et al., 2019; Sipilä et al., 2021; Widmann et al., 2014). The cellular and molecular mechanisms leading to neurocognitive dysfunction in sepsis survivors of are not well understood and further research is needed to develop novel therapeutic strategies to prevent and treat this devastating condition.

Increasing clinical evidence indicates that the endothelium and the blood-brain barrier (BBB) are profoundly affected during sepsis and could play an important role in the development of cognitive impairment (Greene et al., 2024; Gust et al., 2017; Thakur et al., 2021; Widmann et al., 2014). The cerebrovascular endothelium, a key housekeeper of the BBB and neurovascular homeostasis, constitutes a promising therapeutic opportunity. Thus, further examination of the molecular and cellular mechanisms leading to and resulting from BBB and cerebral microvascular dysfunction is required to broaden our understanding of how systemic inflammatory factors signal via the cerebral microvasculature to alter cognitive function and identify novel therapeutic targets. However, these investigations have been hampered by the lack of standardized experimental models to determine the alterations in the cerebral microvasculature and cognition induced by systemic inflammation. To this end, in the present study we have established a mouse model of endotoxemia that recapitulates key aspects of the pathophysiology of sepsis-induced cognitive dysfunction including the induction of an acute systemic hyperinflammatory response, loss of BBB integrity and neurovascular inflammation, followed by alterations in cognitive function in the subacute and chronic phases, after recovery from systemic inflammation. We have validated a scoring system to assess the clinical signs of the systemic inflammatory response. In addition, we have identified key molecular (i.e. induction of the caveolar protein plasmalemma vesicle associated protein, PLVAP, and the antifibrinolytic plasminogen activator inhibitor, PAI-1) and functional alterations in the cerebral microvasculature (i.e. abumin and small molecule BBB leakage and intra/perivascular fibrin/ogen deposition), along with perturbations in neuronal molecular markers associated with cognitive dysfunction of potential clinical relevance, including changes in neurofilament, postsynaptic density protein 95 (PSD-95) and brain-derived neurotrophic factor (BDNF). This work will permit future investigations of the mechanisms underlying systemic inflammation-induced cognitive dysfunction, particularly, the role of the cerebral microvasculature, and provide a reliable model for therapeutic discovery of novel vasoprotective agents in cognitive impairment and other neurodegenerative pathologies.

## MATERIALS AND METHODS

### Animals and LPS treatments-

All animal experiments were approved by the Institutional Animal Care and Use Committee. ∼152 C57/BL6 mice (8-10 weeks, male, Jackson Labs) were used for this study. Animals were housed in stable environmental conditions (23°C, 40% relative humidity and light-dark cycle of 12 hours), with access to food and water *ad-libitum*.

LPS endotoxin (Escherichia coli O111:B4 (Sigma L4391) was prepared as a stock solution at 2 mg/ml in normal saline (0.9% w/v), and further diluted in saline for a 200-250 μl injection per mouse (at a dose of 2 mg/kg) on the day of use. We optimized LPS treatments at 2 mg/kg (intraperitoneally) per mouse body weight using a three-injection routine at 24 hours intervals to induce severe and sustained systemic inflammation. In this model, the mortality rates were ∼10% in first three days following LPS injection (acute phase), and less than 5% in the subacute (days 4-7) and chronic (day 7-1 month) phases. Tissues were harvested immediately following euthanasia and flash-frozen before storage and molecular analyses. (Figure 1).

**Figure 1:**
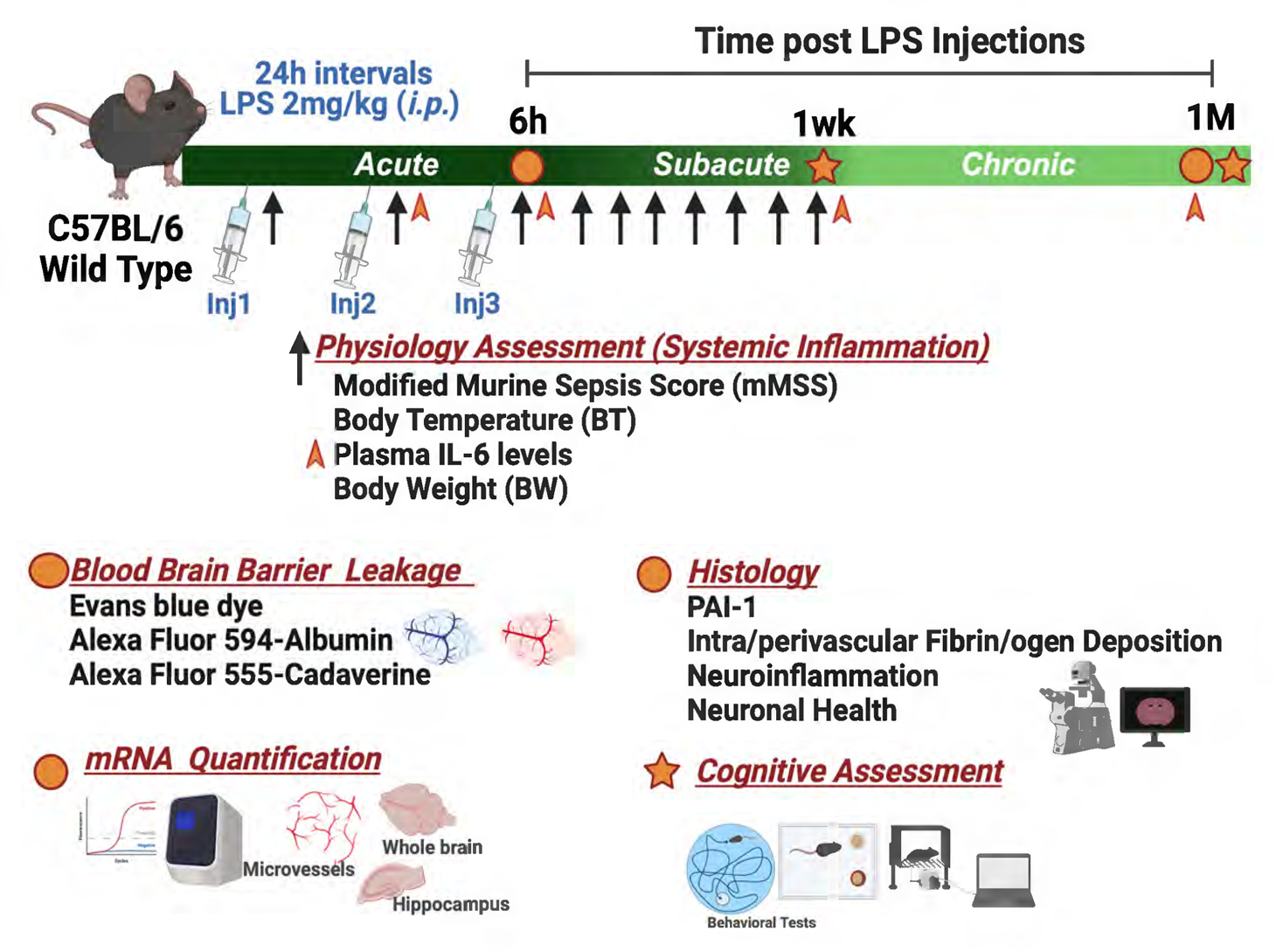
Experimental timeline and subsequent assessments following LPS injections. LPS was administered for three consecutive days. Physiological assessment of the clinical signs of the systemic inflammatory response were conducted 6 hours after each injection and continued daily for one week (black arrows). Blood collection for determination of plasma IL-6 levels was conducted 6 hours after second and third LPS or saline injections, and 1 week and 1 month afterwards (orange arrow heads). Behavioral testing was performed after recovery from the systemic inflammatory response, i.e. one week and one month post LPS injections. Blood-brain barrier (BBB) permeability determination to albumin and small molecules (Evans blue dye [EBD] extravasation assay, histological determination of Alexa Fluor 594-albumin and Alexa Fluor 555-cadaverine leakage), RNA quantification (RT-qPCR analysis), and immunofluorescence imaging were conducted 6h (acute phase) and one month after last LPS injection (chronic phase).

### Physiological assessment of the clinical signs of the systemic inflammatory response-

Mice were assessed every 6h post LPS injection and daily during the first week after LPS administrations to determine murine sepsis scoring, changes in temperature and body weight. Plasma levels of IL-6, a reliable biomarker of systemic inflammation, were also determined. 5-6 mice per group (saline and LPS) were used for these studies.

#### Murine Sepsis Scoring with Nesting Behavior (MSS)

Systemic inflammation was induced by intraperitoneal injection of 2 mg/kg LPS at 24h intervals over three days (Figure 1). To assess the clinical signs of systemic inflammation we designed a multi-parametric assessment scale that combines the murine sepsis score (MSS) (Shrum et al., 2014) and nesting behavior (Deacon, 2006) rating scale (modified MSS, mMSS). The assessed parameters included features such as appearance (piloerection, puffiness), level of consciousness (posture impairment, social engagement), activity (exploratory behavior, movement speed), response to stimulus (reaction time to sound and touch stimuli), eye condition (presence of secretions or closed eyelids) respiratory rate (presence of apneas, slowed breathing pace) and nesting behavior, which are affected following inflammation and are reminiscent of the sepsis phenotype. For the nesting assessment, nesting pads (Nestlets, Ancare Corp., Bellmore NY) were placed in the mice cages after every saline or LPS injection. Healthy mice quickly tear the pads to smithereens and build a nest from scratch, whereas septic mice ability to build the nest decreases, leading to larger portions or even intact nesting pads). A new nesting pad was placed every day. This scoring system is described in detail in Table 1.

**Table 1.** Assessment Parameters of modified Murine Sepsis Score (mMSS)

| Score | 0 | 1 | 2 | 3 | 4 |
| --- | --- | --- | --- | --- | --- |
| Appearance | Smooth | Patches of hair piloerected | Majority of back is piloerected | mouse appears “puffy” | Mouse appears emaciated |
| Level of Consciousness | Active | Still active, but avoids standing upright | Slowed, still ambulant | Impaired, only provoked move with a tremor | Severely impaired, stationary with possible tremor |
| Activity | Normal | Slightly suppressed moving around bottom of cage | Suppressed Stationary with occasional investigation | No activity | No activity and tremors, particularly in the hind legs |
| Response to stimulus | Immediately to auditory stimulus or touch | auditory: slow or no, touch: strong (moves to escape) | auditory: no, touch: moderate (moves a few steps) | auditory: no, touch: mild (no locomotion) | auditory: no, touch: little or no |
| Eyes | Open | Eyes not fully open, possibly with secretions | Eyes at least half closed, possibly with secretions | Eyes half closed or more, possibly with secretions | Eyes closed or milky |
| Respiration rate | Normal, rapid, not quantifiable by the eye | Slightly decreased (not quantifiable by the eye) | Moderately reduced (quantifiable by the eye) | Severely reduced (0.5 s between breaths) | Extremely reduced (>1 s between breaths) |
| Nesting behavior | Normal, fully scratched and nested | Majority remains, fluffy with some parts intact | Small piece of nest, partially fluffy | Large portion remains intact | Largely untouched |

#### Changes in temperature and body weight

Body weight measurements were recorded using a portable weighing balance. Body temperature was recorded using a rectal thermal probe.

#### Plasma levels of IL-6

Blood was obtained via submandibular vein puncture using lancets and collected into 0.5 M EDTA-coated capillaries, 6 hours after second and third LPS or saline injections, 1 week and 1 month after last LPS or saline injection. Plasma was then obtained by centrifugation of the collected samples for 15 minutes at 1500 x g, and stored at −20°C until further analyses. Plasma IL-6 levels were then determined using a commercially available mouse IL-6 Quantikine ELISA Kit (R&D Systems, M6000B). 5-6 mice per group (saline and LPS) were used for plasma collection.

### Determination of BBB leakage-

#### Evans blue dye (EBD) extravasation assay

*BBB leakage* to albumin was assessed by EBD extravasation assay in saline and LPS-injected mice at 6 h and 1 month after third injection, as previously described (Kim et al., 2015). 5-8 animals per group were used for these studies. 2% EBD (wt/vol in PBS) at 0.2 mg/g was injected into the right jugular vein and allowed to circulate for 2 hours. Under deep anesthesia, mice underwent transcardial perfusion with PBS (+5 mM EDTA) via a 23-gauge butterfly needle through the left ventricle until a clear fluid was obtained from the right atrium (∼20 mL). The brains were removed, and the olfactory bulb and cerebellum were cut off. The resulting brain hemispheres were cut in half with half taken for EBD measurement and the other half for RNA extraction. The hemispheres were weighed, and EBD was extracted with 50% trichloroacetic acid (TCA) as previously described (Zhang et al., 2013), (Yanagida et al., 2017), (Kim et al., 2015). Briefly, the one brain hemisphere was weighed and then homogenized in 1 ml of 50% TCA (weight/volume) and subjected to centrifugation at 15, 000 g for 15 min. This centrifugation was repeated an additional time to remove the leftover debris. Concentration of EBD in the supernatant was measured in triplicates using fluorescence spectroscopy (620_ex_/680_em_).

#### Histological determination of Albumin and Cadaverine BBB leakage

Albumin and small molecule (Cadaverine) BBB leakage were determined in separate groups of mice, treated with saline or LPS, at 6 h or 1 month after third injection (3-4 animals per group). Alexa Fluor 594 Albumin (5mg/mL, Thermo Fisher) solution in saline was injected into the right jugular vein. After allowing to circulate for 2h, mice were transcardially perfused as previously described with cold PBS/EDTA followed by 4% PFA and further processed for histology. Likewise, 1.5 mg/mL Alexa Fluor® 555 Cadaverine was injected at a 6 µg/g dose using the same approach.

#### Cerebral Microvessel isolation-

Cerebral microvessels were isolated from saline and LPS-injected mice at 6 h and 1 month after third injection, as previously described (Lee et al., 2019). 5 animals per group were used for microvessel isolation. To minimize cell activation, all procedures were conducted in a cold room. Ipsilateral cortices were homogenized with MCDB131 medium (Thermo Fisher Scientific, 10372019) with 0.5% fatty acid free BSA (Millipore Sigma, 126609). The homogenate was centrifuged at 2000 g for 5 minutes at 4°C. The pellet was suspended in 15% dextran (molecular weight ∼70 kDa, Millipore Sigma, 31390) in PBS and centrifuged at 10000 g for 15 minutes at 4°C. The pellet containing the microvessels was resuspended in MCDB131 with 0.5% fatty acid free BSA and centrifuged at 2000 g for 10 min at 4°C.

#### mRNA quantification, cDNA synthesis and quantitative PCR (qPCR)-

6 h or 1 month after third LPS or saline injections, RNA was extracted from whole brain (3-5 animals per group), isolated cerebral microvessels (5 animals per group) or hippocampus (3-5 animals per group) using Trizol (ThermoFisher, MA) and the Qiagen RNeasy Mini Kit (Qiagen, MD) as previously described (Lee et al., 2019). cDNA was produced using Verso-Reverse Transcriptase (RT) kit with Random Hexamer Primers (ThermoFisher, CA) and diluted with nuclease free water. SYBR Green with Rox dye (Quanta Bio, MA)-based qPCR was performed to determine the relative expression levels of target genes, Claudin-5 (Cldn5), Plvap, Caveolin-1 (Cav1), Serpine1 and Bdnf. qPCR reactions were run in duplicates or triplicates on a 96 well plate loaded on to an ABI-7500 Sequence Detection System PCR machine (Applied Biosystems, CA) and the averages of the duplicates/triplicates Ct values were calculated. Ct values of target genes were normalized to expression levels (Ct values) of hypoxanthine phosphoribosyltransferase (Hprt) RNA, as ΔCt values. For saline and LPS samples, the relative gene expression levels were expressed as ΔΔ Ct values by subtracting the average ΔCt values of the saline group from each ΔCt value. Fold change in target gene expression was calculated by comparing the 2^-ΔΔ^ ^Ct^ values of the LPS samples and saline samples. The primer sequences and gene ID numbers were the following:

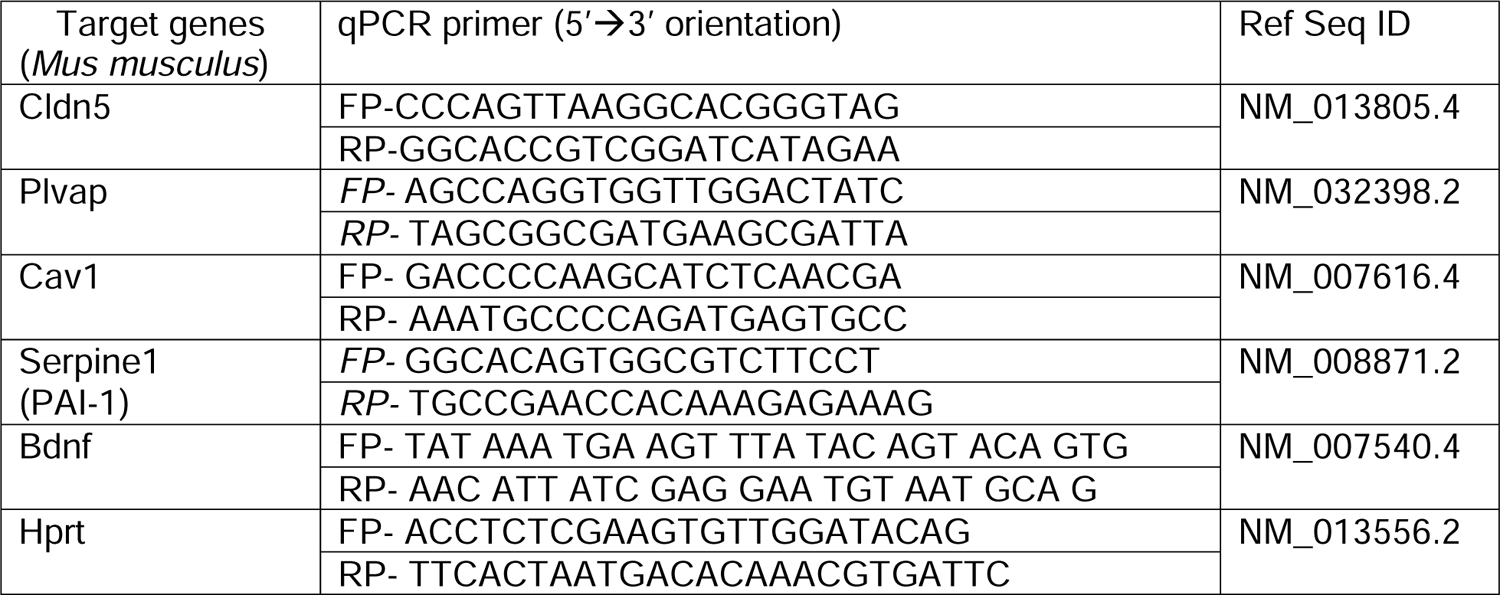

#### Barnes Circular Maze-

1 week after 3^rd^ LPS or saline injection, mice were subjected to Barnes maze test. 9-10 animals/group were used. The Barnes circular maze consisted of a circular open surface (90 cm in diameter) elevated to 90 cm by four wooden legs (O’Leary et al., 2013), (Faraco et al., 2019). There were 20 circular holes (5 cm in diameter) equally spaced around the perimeter and positioned 2.5 cm from the edge of the maze. No wall and no intra-maze visual cues were placed around the edge. A wooden plastic escape box (11×6×5 cm) was positioned beneath one of the holes. Three neon lamps and a buzzer were used as aversive stimuli. The Any-Maze tracking system (Stoelting) was used to record the movement of mice on the maze. Extra-maze visual cues consisted of objects within the room (table, computer, sink, door) and the experimenter. Mice were tested in groups of seven to ten, and between trials they were placed into cages, which were placed in a dark room adjacent to the test room for the inter-trial interval (30–60 minutes). No habituation trial was performed. The acquisition phase consisted of 3 consecutive training days with three trials per day with the escape hole located at the same location across trials and days. On each trial a mouse was placed into a start tube located in the center of the maze, the start tube was raised, and the buzzer (white noise) was turned on until the mouse entered the escape hole. After each trial, mice remained in the escape box for 60 seconds before being returned to their cage. Between trials the maze floor was cleaned with 10% ethanol in water to minimize olfactory cues. For each trial mice were given 3 minutes to locate the escape hole, after which they were guided to the escape hole or placed directly into the escape box if they failed to enter the escape hole. Parameters recorded for learning performance were below: (1) the latency to locate (primary latency) and (2) the distance traveled before locating the escape hole (Faraco et al., 2019). On days 4, the location of the escape hole was moved 180° from its previous location and two trials per day were performed. Any Maze v5.3 was used for collection and analysis of the behavioral data.

#### Novel object recognition (NOR)-

1 month after 3^rd^ LPS or saline injection, mice were subjected to NOR test (12-14 per group). The NOR task was conducted under dim light in a plastic box with dimensions 29 cm × 47 cm × 30 cm high. Stimuli consisted of plastic objects that varied in color and shape but had similar size (Grayson et al., 2015); (Cohen et al., 2015). A video camera mounted on the wall directly above the box was used to record the testing session for offline analysis. Mice were habituated to the testing environment and chamber for 24 hours prior to testing. Thereafter, mice were placed in the same box in the presence of two identical sample objects and were allowed to explore for 5 minutes. After an intersession interval of 24 hours, one of the two objects was replaced by a novel object, and mice were returned to the same box and allowed to explore for 5 minutes (Faraco et al., 2019). Exploratory behavior was then assessed manually by a researcher blinded to the treatment group. Exploration of an object was defined as the mouse sniffing the object or touching the object while looking at it. Placing the forepaws on the objects was considered as exploratory behavior but climbing on the objects was not. Discrimination ratio was calculated by dividing the difference of the amount of exploration time between novel and familiar object by the total exploration time. Any Maze v5.3 was used for collection and analysis of the behavioral data.

#### Associative Fear Conditioning-

A battery of behavioral protocols was carried out to assess fear response 1 month after 3^rd^ LPS or saline injection (12-29 mice/group). Fear test was carried out using established protocols (Meyer et al., 2019).

Protocols were carried out in standard conditioning chambers (Med Associates). The chambers (30 × 24 × 27 cm) consisted of clear acrylic front and back walls, aluminum sides, stainless steel grid floors, and a clear acrylic top. Each chamber was outfitted with a speaker located 13 cm above the grid floor, used to present the auditory conditioned stimulus. Delivery of a foot shock through the grid floor served as the aversive unconditioned stimulus. For context setup, a chamber with LED Stimulus Light (50 lux) mounted 18 cm above the grid floor to provide background illumination containing olfactory cues (peppermint, 1/1000 in ethanol) was used. Each chamber was enclosed in a sound-attenuating cubicle (71 × 59 × 56 cm) with a 28 V DC exhaust fan to provide airflow and background noise. Video Freeze® software (Med Associates) was used to control experiments, and all trials were videotaped for analysis. Freezing, defined as the absence of visible movement except that required for respiration, was quantified using the Video Freeze® software set at a motion threshold of 18 Units for automatic scoring.

#### Fear Conditioning

Mice were acclimated to the conditioning chamber for 2 minutes prior to five tone presentations (5 kHz, 80 dB, 20 s duration) that co-terminated with a foot shock (0.5 mA, 1 s duration). The intertrial interval (ITI) between the tone-shock pairings was set at 1 minute. Mice remained in the conditioning chamber for 1 minute after the final tone-shock pairing before being returned to their home cages. The baseline motion was recorded during acclimation of first 2 minutes, then the percentage time spent freezing during each tone (20 s) was calculated, providing an index of cued fear learning.

#### Contextual Fear Recall Test

Mice were tested for contextual fear 24 hours after fear conditioning. Mice were returned to the context setup for 5.5 minutes during which no tones or shocks were presented. Upon completion of the contextual fear test mice were returned to their home cages. The percentage of time spent freezing across the first 2 minutes of entire 5.5 minutes session was calculated, providing an index of contextual fear memory.

#### Immunofluorescence-

6 h or 1 month after third LPS or saline injections, mice were perfused with cold PBS under deep anesthesia and subsequently with 4% PFA in PBS solution. 3-5 mice per group were used for these histological analyses. The brains were removed, postfixed with 4% PFA for 24 h, transferred to 30% sucrose solution in PBS, embedded in optimal cutting temperature (OCT) compound, frozen on liquid nitrogen and placed in a − 80 C freezer for storage. Ten μm thick sections were cut on a cryostat (Leica Microsystems, Buffalo Grove, IL) through +0.5 to −2.6 mm from Bregma. Sections were washed three times with PBS and then blocked with blocking solution (5 % bovine serum albumin, 0.8 % skim milk, and 0.3 % Triton X-100 in TBS) for 1 h and incubated with the specified primary antibodies in blocking solution overnight at 4 °C, followed by the appropriate secondary antibodies and 4’, 6-diamidino-2-phenylindole (DAPI) for 1 hours at room temperature and were mounted onto slides. The cerebral microvasculature was identified by immunofluorescence for the endothelial-specific marker, glucose transporter 1 (Glut1), as previously described (Callegari et al., 2023). The following antibodies were used in this study: anti-Glut1, Alexa Fluor 488 conjugated (Rabbit polyclonal, from Millipore, 1:100), anti-Serpine1 (Rabbit polyclonal, from Innovative Research, 1:100), anti-Ionized calcium binding adaptor molecule1 (Iba1, Rabbit polyclonal, from Wako, 1:100), anti-Glial fibrillary acidic protein (Gfap, Rabbit polyclonal, from Abcam, 1:100), anti-Bdnf (Rabbit monoclonal, from Abcam, 1:100, which recognizes pro-BDNF and biologically active BDNF isoforms), anti-hyperphosphorylated neurofilament heavy chain (p-NF, mouse monoclonal, clone SMI 31P, from Biolegend, 1:100), Purified anti-Neurofilament H (NF-H), Nonphosphorylated Antibody, clone SMI32 (801701, Biolegend, 1:100), Recombinant Alexa Fluor 647 Anti-P2Y12 antibody ([EPR26298-93] ab308432, Abcam), anti-Fibrinogen (Rabbit Polyclonal ab34269, Fisher Sci), Recombinant Anti-PSD-95 [EPR23124-118] (ab238135, Abcam).

#### Image acquisition and analysis-

Images were acquired on an Agilent BioTek Lionheart FX Automated microscope (20x Magnification Objective) (Agilent, US) and quantified using BioTek Gen5 software for pNF (SMI31), SMI32, P2Y12/IBA1, PSD95 and fibrinogen, and QuPath software for PAI1, GFAP, BDNF. Images for Alexa Fluor 594-Albumin and Alexa Fluor 555-cadaverine analysis were acquired using a Leica Stellaris Confocal Microscope (40x Magnification Objective) and analyzed using QuPath software. For all analysis, images were acquired from 2 sections per mouse brain, one anterior (+0.5 to −0.3 mm from Bregma) and one posterior (between −1.8 to −2.6 mm from Bregma). A total of 6-10 brain sections from 3-5 mice were quantified. For analysis using BioTek Gen5, 10-12 regions of interest (ROI) from cortex and 3-6 ROI from hippocampus per mouse brain were acquired and quantified. For analysis using QuPath, whole brain images were acquired and 10-12 (ROI) from cortex and 18-20 small ROI (4-6 ROI for BDNF) from hippocampus were quantified per mouse brain (Extended Data Figure 3-2). Vessel area on each ROI was automatically detected using GLUT1 (endothelial-specific marker) staining and QuPath’s built-in cell segmentation algorithms as previously described (Callegari et al., 2023). Albumin puncta were quantified within the vessel and in a 10 µm region around the vessel, and normalized by vessel area. PAI-1 total intensity in vessel detected area, fibrinogen positive area in vessel detected area, or IBA1, P2Y12, GFAP, BDNF, PSD-95, SMI31, SMI32, and Cadaverine positive area detected by thresholding on each ROI were quantified and normalized by the mean value of the ROI of saline injected mice. In P2Y12+ regions, the IBA1+ integrated density was used as a thresholder to detect P2Y12+/IBA1+ regions. All quantified data was exported to Excel and Prism9 for further analysis.

#### Statistical analysis-

For EBD leakage assay, RT-qPCR, image analyses quantification and behavior assessment statistical analyses were conducted using GraphPad Prism9 software. All values reported are mean ± SEM.

For comparisons between two groups, an Unpaired t-test was used. For data involving time series, a two-way analysis of variance (ANOVA) followed by Sidak’s multiple comparison test was used. For comparisons among the 3 groups (control group, 6 hours after the final LPS administration, and one month after), a one-way ANOVA followed by Tukey’s multiple comparison test was used. P<0.05 was defined as statistically significant. (*p<0.05, **p<0.005, ***p<0.0005, ****p<0.0001). A statistical table for all analyses is provided in Extended Data Table 1.

## RESULTS

### Characterization of the endotoxemia-induced mouse model of systemic inflammation

We established a mouse model of endotoxemia to recapitulate important clinical and neurological features of systemic inflammatory response syndrome and sepsis, including the induction of a hyper-inflammatory response *via* the activation of pathogen recognition receptors, as well as key aspects of the pathophysiology of cognitive dysfunction induced by severe systemic inflammation including BBB leakage, neuroinflammation and memory impairment (Figure 1). We administered LPS treatments at 2mg/kg per mouse body weight using a three I.P. injection protocol at 24h intervals to induce a sustained systemic inflammatory response (Figure 2). We next determined the clinical signs of systemic inflammation using a multi-parametric assessment scale, i.e. murine sepsis scores, MSS, and changes in body temperature. Our modified MSS (mMSS) included a previously described scoring system (Shrum et al., 2014) and a nesting behavior score (Deacon, 2006). Mice were scored on six variables including grooming, activity, consciousness, eye secretions, respiratory rate, stimulus response and nesting behavior (Table 1). LPS administration caused a rapid and acute response in dosed mice, as reflected by an immediate increase in mMSS following the initial injection. The mMSS continued to increase with every injection peaking after the 3^rd^ dose and returned to normal baseline 72-96h post-final injection (Figure 2A). Changes in the individual parameters are shown in Figure 2-1. Regarding body temperature, LPS-dosed mice showed a 0.5 °C decrease in body temperature after the second injection. At 24h following the third LPS administration, dosed mice showed a significant reduction (33.0 °C) in body temperature compared with the saline group (34.0 °C). In a similar pattern to mMSS, animals’ body temperature slowly recovered beyond 24h and reached baseline levels at 72-96h (Figure 2B). These changes in mMSS and body temperature suggest that the acute systemic inflammatory response induced by LPS resolved near 72h after last injection. We also confirmed the development and resolution of the systemic inflammatory response by measuring plasma levels of IL-6, a well-established biomarker of systemic inflammation and sepsis morbidity (Remick et al., 2002; Shapiro et al., 2010). IL-6 plasma levels sharply increased after LPS injections and returned to baseline levels by 1 week (Figure 2C). Finally, LPS injection also caused a marked reduction in body weight following the initial dose, and steadily decreased with subsequent administrations (Figure 2D). At 24h post-LPS injection, mice showed a 18% reduction in body weight compared with the control group. The animals’ weight consistently increased beyond 24h post-administration and reached pre-treatment weight 1 week after LPS dosing. Conversely, saline-treated animals showed no differences in mMSS, body temperature, plasma IL-6 levels or weight throughout the different timepoints. Mortality rates were less than 10% in first 3 days during LPS injection (acute phase) and less than 5% in the subacute (days 4-7) and chronic (day 7-1 month) phases and 0% in saline-injected mice.

**Figure 2:**
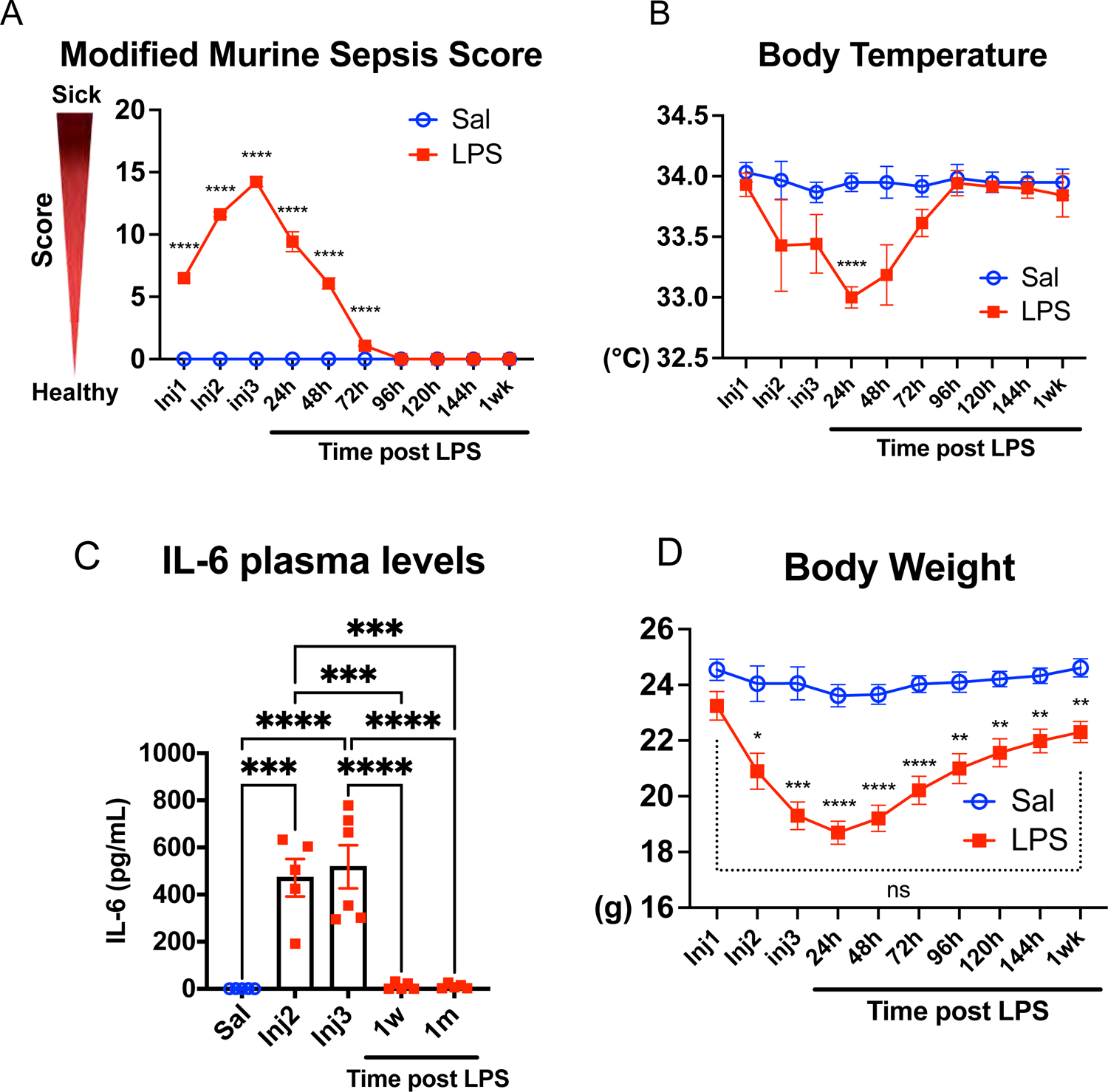
LPS injections induce an acute systemic inflammatory response that resolves within 72-96 hours. Following LPS injections, assessments of the murine sepsis score (MSS), body temperature, IL-6 plasma levels and body weight, were conducted to evaluate physiological changes and gauge the severity of systemic inflammation. (A) The MSS score surged immediately following the initial injection, reaching its peak after the 3rd dose. Following LPS treatment, the MSS score rapidly diminished, matching the saline control levels from 96 hours onwards. (B) Body temperature exhibited a decline during LPS treatments and returned to baseline levels 96 hours after the last LPS dose. (C) IL-6 plasma levels (pg/mL) were significantly increased following LPS injection and returned to baseline levels at 1 week after LPS administration (N=5-6/group). (D) Body weight demonstrated a pronounced decrease during the LPS injection period, began recovery 24 hours after the final LPS administration, and nearly reached baseline one week post the last injection. The mean of individual value±SEM are shown. A, B and D: N=7 Saline, N=13 LPS. *p<0.05, **p<0.005, ***p<0.0005, ****p<0.0001, two-way ANOVA followed by Sidak’s multiple comparison test.

Altogether the mMSS, body temperature and IL-6 data indicate that the described model of endotoxemia induced an acute systemic hyper-inflammatory response immediate after the first LPS injection, reaching a peak after the 3^rd^ dose and resolving at near 72 hours after last injection.

### Acute and chronic functional and molecular alterations in the cerebral microvasculature and neuroinflammation in the endotoxemia-induced systemic inflammation model

In order to study the impact of systemic inflammation on BBB function we used the EBD assay to determine albumin leakage into the brain parenchyma. We observed a robust increase in EBD extravasation in whole brain homogenates, in the acute phase of systemic inflammation (3.7±0.129 fold, 6 h following last LPS injection) which returned to near baseline levels at 1 month (Figure 3A). To confirm these data histologically and to determine the permeability of macromolecules and small molecules through the BBB in our model, we used two intravascular tracers, Alexa Fluor 594-albumin (∼66kDa) and Alexa Fluor 555-cadaverine (∼1kDa), as previously described (Armulik et al., 2010; Knowland et al., 2014; Yanagida et al., 2017). Consistent with the EBD extravasation assay, we observed a robust BBB leakage to Alexa Fluor 594-albumin in the acute phase both in hippocampus (Figure 3 B, C, 3.01±0.48 fold) and cortex (Figure 3-1 A, B, 2.7±0.23 fold), but not at 1 month. Remarkably, small molecule (Alexa Fluor 555-cadaverine) leakage was robust in the acute phase and persistent in the chronic phase both in hippocampus (Figure 3D-E, 6.3±0.8 and 3.9±0.56 fold, respectively) and cortex (Figure 3-1 C, D). We also analyzed the mRNA levels of molecules governing microvascular function in cerebral microvessels isolated from both control and LPS-treated mice, such as tight junction proteins (Cldn5), molecules governing vesicular trafficking (PLVAP, Cav1) and coagulation (plasminogen activator inhibitor-1, PAI-1, encoded by the transcript Serpine-1). LPS-induced inflammation caused a marked reduction in Cldn5 (∼62% decrease) mRNA levels at 6h, along with a significant increase in the expression of Plvap (7.8±1.1 fold) and the anti-fibrinolytic PAI-1 (Serpine-1 transcript, 3±0.58 fold). Interestingly, while the alterations in transcripts governing barrier function reverted to near baseline levels in the chronic phase (1 month after induction of systemic inflammation), the increase in procoagulant Serpine-1 transcript levels was still present at 1 month (Figure 3F, 2.8±0.37 fold). Moreover, the histopathological immunofluorescence (IF) analysis of hippocampal (Figure 3G-H) and cortical (Figure 3-1 E, F) microvessels using Glut-1 antibody as an endothelial-specific marker confirmed the robust increase in the number of PAI-1+ vessels both at 6h and 1 month (86% and 32% increase, respectively) following LPS administration compared to saline. Since increased levels of PAI-1 contribute to impaired fibrin degradation, fibrin deposition and microvascular dysfunction in sepsis (Shapiro et al., 2010), we also determined the functional consequences of this sustained PAI-1 upregulation in the cerebral microvasculature by quantifying the levels of fibrin in the cerebral microvasculature in our model. IF analyses with anti-fibrin/ogen and Glut-1 antibodies revealed increased intra/perivascular fibrin/ogen deposition in the chronic phase both in hippocampus and cortex (1.7± 0.1 and 1.8± 0.3-fold, respectively, Figure 3-1 G-J). Altogether, these data indicate that the established endotoxemia model induces cerebral microvascular inflammation (assessed by BBB permeability), expression of procoagulant molecules and intra/perivascular fibrin/ogen deposition, both in the acute phase and 1 month after recovery from systemic inflammation.

**Figure 3:**
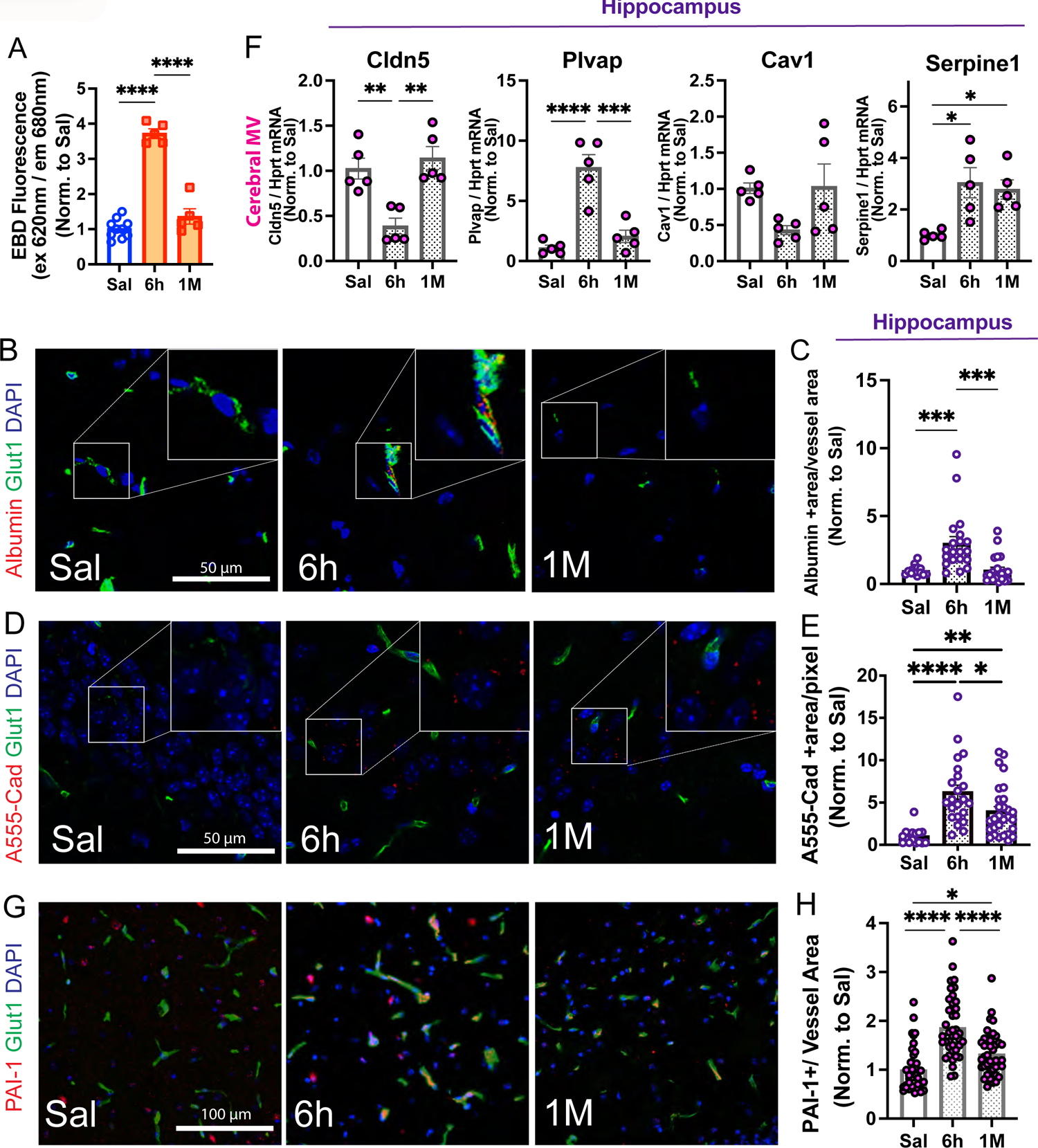
LPS induces acute and chronic alterations in microvascular inflammation and integrity. (A) Blood-brain barrier (BBB) permeability, indicated by Evans blue dye (EBD) leakage (albumin), was evident at 6h post LPS administration and returned to baseline levels by 1 month after the final LPS injection. Evans blue dye (EBD) fluorescence in brain homogenates normalized to saline is shown. N=5-10 mice/group. (B-E) Histopathological validation of BBB leakage in hippocampus. N=5-8/group. (B, C) Albumin leakage in hippocampus using Alexa Fluor 594-albumin as a tracer shows an increase in albumin+ area at 6h following LPS administration that returns to baseline levels at 1 month. (D, E) Alexa Fluor 555-cadaverine leakage showed an increase in BBB permeability in hippocampus 6h after LPS treatment that remained elevated at 1 month. Panels B and D show representative pictures (40x magnification) of the hippocampus with a 2x digital zoom inset. Scale bar=50μm (F) LPS-induced systemic inflammation caused a marked reduction in Cldn5 together with an increase in Plvap and the anti-fibrinolytic Serpine-1 mRNA (which encodes PAI-1), assessed by RT-qPCR in isolated cerebral microvessels (MV). Notice that Serpine-1 mRNA levels remained elevated in the chronic phase (1 month). Fold change LPS compared to saline is plotted. Data points represent the average of duplicates or triplicates of each mouse. Individual values and average ± SEM are shown. N=4-5 mice/group. (G, H) Histopathological validation and quantification of PAI-1 alterations (red channel) in the cerebral microvasculature (Glut1 positive areas, green channel) in the hippocampus. N=3 mice/group. Panel G shows representative pictures of the hippocampus (20x magnification). Scale bar=100μm. Nuclear staining (DAPI) is shown in the blue channel in all images. Each data point represents one mouse brain in panels (A) and (F), and one region of interest (ROI) from the entire scanned image in panels C, E and H. The individual values and the mean±SEM are shown. All data were normalized by the mean of Saline. *p<0.05, **p<0.005, ***p<0.0005, ****p<0.0001, one-way ANOVA followed by Tukey’s test.

Next, we quantitatively assessed the levels of neuroinflammatory markers in the cerebral parenchyma following systemic inflammation through IF analyses. We focused on the expression dynamics of Gfap, a canonical indicator for activated astrocytes, the ADP receptor P2Y12, as a marker for homeostatic microglia (Krasemann et al., 2017; Mildner et al., 2017) and Iba1, an indicator of activated microglia/macrophages. These markers were evaluated in two cerebral regions: the cortex and the hippocampus. Data demonstrates a pronounced increase in GFAP (Figure 4A, B, 1.4±0.06 fold) and IBA1 immunoreactivity in hippocampus (2.28±0.44-fold, Figure 4C, D), together with a decrease in the homoeostatic microglial marker P2Y12 (22% decrease, Figure 4E, F), 6 hours post-final LPS administration, underscoring an immediate neuroinflammatory response during the acute phase. Analysis of P2Y12+ and IBA1+ (double positive) immunoreactivity suggested an increase in activated microglia (2.8±0.6-fold, Figure 4G-H). One month after recovery from systemic inflammation, quantification of GFAP and IBA1 alterations revealed a regression of these heightened levels (Figure 4A-D). Interestingly, P2Y12 immunopositivity at 1 month remained significantly lower (20%) compared to saline mice, which together with the trend in increased IBA1+/P2Y12+ immunoreactivity (58% higher, p=0.002 by t-test, 1 month vs saline), suggests continued microglial activation in the chronic phase. Analyses of the alterations of these markers in cortex also followed a similar trend (Figure 4-1), suggesting that astrocyte activation surged during the acute phase, then returned to normal levels in the chronic phase, while microglial activation remained elevated. These results suggest a temporal pattern of neuroinflammatory marker expression post-systemic inflammation, with an initial spike in the acute phase and a significant decrease in the chronic phase with lingering, low-grade microglia activation.

**Figure 4:**
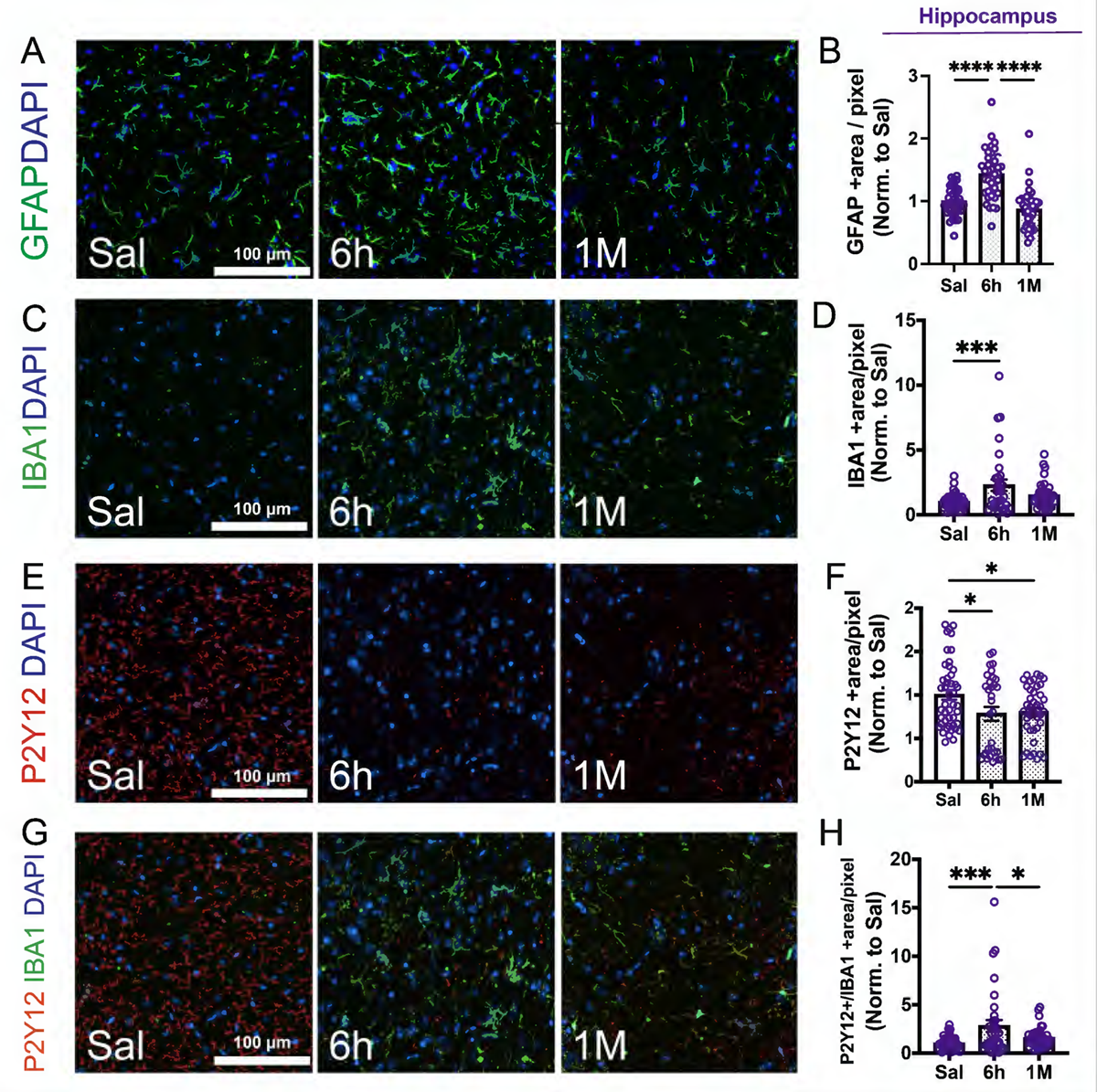
Endotoxin-induced systemic inflammation leads to increased expression of neuroinflammatory markers. Immunofluorescence analyses and quantification for (A, B) Glial fibrillary acidic protein (GFAP, green channel), (C, D) Ionized calcium-binding adaptor molecule 1 (IBA1, activated microglia/macrophage marker, green channel), (E, F) ADP receptor P2Y12 (P2Y12, homeostatic microglia marker, red channel) and (G, H) P2Y12/IBA1 (merged) staining in the hippocampus from Saline-treated controls, and 6 hours and 1 month post LPS administration. (B, D, F, H) Quantitative analysis representing the total area of GFAP+, IBA1+, P2Y12+ and IBA1+P2Y12+ staining in hippocampus. Notice the transient increase in GFAP and IBA1 immunopositivity at 6h and the decrease in P2Y12. All nuclei were labeled with DAPI (blue channel). Panels A, C, E and G show representative pictures (20x magnification). Scale bar=100μm. Each data point symbolizes one region of interest (ROI) from the entire scanned image normalized by the mean of Saline. N=3-5 mice per group. The individual values and the mean±SEM are shown. ****p<0.0001, one-way ANOVA followed by Tukey’s test.

In summary, our data indicate that the described LPS model of severe systemic inflammation causes both immediate and lasting molecular and functional alterations in the cerebral microvasculature, as well as neuroinflammation. Key changes in the acute phase include a significant increase in BBB permeability, the acquisition of a pro-coagulant phenotype in the cerebral microvasculature, and elevated expression of neuroinflammatory markers. This acute neurovascular inflammatory response tends to decrease 1 month after recovery from systemic inflammation. However, BBB permeability to small molecules, heightened PAI-1 expression in the cerebral microvasculature, and intra/perivascular fibrin/ogen deposition were sustained in the chronic phase, alongside persistent microglial activation.

### Impairment in spatial, recognition and contextual fear memory in the endotoxemia-induced systemic inflammation mouse model

Given the alterations in neurovascular inflammatory markers observed after endotoxemia, particularly in the hippocampus, next we aimed to determine the impact of systemic inflammation in memory and cognition. We performed a series of behavioral tests largely dependent on hippocampal integrity (Bach et al., 1995; Chen et al., 2011; Cohen et al., 2015) to assess memory after recovery from endotoxemia, at day 7 and 1 month, when mice had recovered from systemic inflammation, and had no visible signs of locomotive or physiological deficits as assessed by the mMSS and plasma IL-6 levels. We performed spatial learning and memory assessment at 1-week post-systemic inflammation using the well-established Barnes circular maze paradigm (Figure 5A-D), the earliest time point when the mice had no visible sign of sickness or locomotive deficits (Figure 2). Mice were subjected to three consecutive training days with three trials per day, with the escape hole located at the same location across trials and days and to two additional trials on day 4^th^, in which the scape hole was moved to 180° from its previous location. Both saline and LPS-injected groups exhibited similar locomotive activity with a decrease in the traveled distance from day 1 to a minimum on day 3, showing no significant differences between them (Figure 5B). From day 1 to day 3 of test, both groups showed decreasing time of primary latency, which means both groups learned the location of escape hole to the same degree (Figure 5C). Interestingly, when the escape hole was repositioned by 180° on day 4, as shown in Figure 5A, LPS-treated mice required more than twice the time (∼2.3-fold more, p<0.05) to locate the escape hole when compared with saline-treated mice (Figure 5C, D), indicating that they had impaired spatial memory.

**Figure 5:**
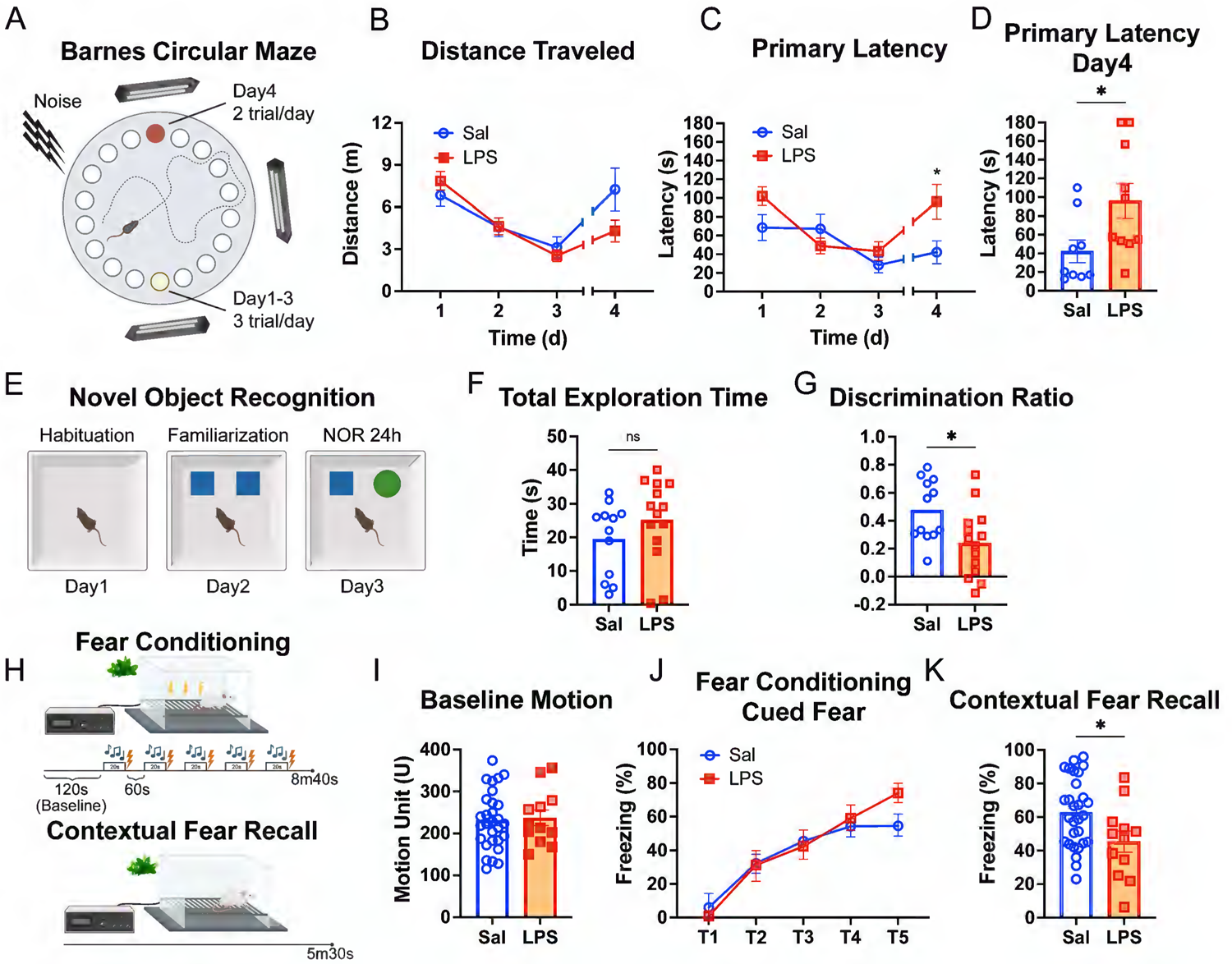
Deficits in spatial, recognition and contextual fear memory and learning after recovery from systemic inflammation. (A) Schematic representation of the Barnes Circular Maze (BCM) procedures conducted one week post recovery from systemic inflammation. (B) Saline and LPS-injected mice showed similar exploratory behavior assessed by the distance traveled. (C, D) LPS-treated mice exhibited spatial memory deficits compared to saline injected mice. They required more time to locate the escape hole on Day 4 following its relocation (primary latency). (D) Individual primary latency values at day 4 are shown. *p<0.05, Two-way ANOVA followed by Sidak’s test or Unpaired t-test. (E) Schematic representation of the Novel Object Recognition (NOR) procedures. One month post systemic inflammation, (F) Saline and LPS-injected mice showed equivalent exploratory movement, assessed by the total exploration time. (G) LPS-treated mice show recognition memory deficits compared to saline, failing to recognize the novel object, determined by the lower discrimination ratio. *p<0.05, Unpaired t-test. (H) Schematic representation of the Fear Conditioning and Contextual Fear Test procedures, conducted 1 month after recovery from systemic inflammation. (I) Baseline activity, determined by motion unit automatically calculated by software, prior to the application of electric shock was comparable between both groups. (J) LPS-treated mice did not exhibit deficits in fear learning, assessed by the percentage of freezing time after the tone. (K) LPS-treated mice displayed impairments in contextual fear recall compared to saline injected mice, assessed by the percentage of freezing time after being in the same context but without the tone or application of the electric shock. *p<0.05, Unpaired t-test (I, K) or two-way ANOVA followed by Sidak’s test (J).

Next, we conducted the novel objection recognition (NOR) test to understand whether our model of severe systemic inflammation impacts the recognition memory in mice at 1-month after recovery (chronic phase, Figure 5E). On average, there were no significant differences between groups in the total amount of exploration time (Figure 5F). However, LPS-treated mice showed significantly lower discrimination ratio between novel object and familiar object (∼25%, p<0.05) compared to the saline control group at 24h post familiarization (Figure 5G), indicating that systemic inflammation negatively impacted the ability to remember and recognize the novel object.

To understand the lasting effects of systemic inflammation on emotional memory, we studied mouse performance in fear conditioning and contextual fear recall at 1 month post LPS administration (Figure 5H). On day 1, both saline and LPS groups were subjected to fear conditioning by tone test. No differences in baseline motion or percentage of freezing time after each tone were found between groups (Figure 5I, J), indicating that fear learning was not impacted by systemic inflammation. However, when on day 2, mice from both saline and LPS groups were exposed to the context in which they received the adverse stimulus the day before (LED light and smell of peppermint in the paradigm), LPS-treated mice froze significantly less (∼25%, p<0.05) than the saline group during the first 2 minutes of contextual test (Figure 5K). These data indicate that contextual fear memory, which is primarily dependent on hippocampal function (Chen et al., 2011), was impaired in mice 1 month after the recovery from systemic inflammation compared to the saline-injected group.

Altogether, these data indicate that the established bacterial endotoxemia model triggers a transient state of systemic inflammation and a persistent learning and memory impairment after recovery.

### Alterations in neurofilament, post-synaptic density protein-95 and brain-derived neurotrophic factor expression in the chronic phase after recovery from systemic inflammation

Given the deficits in spatial, recognition and contextual fear memory observed after recovery from systemic inflammation, we aimed to identify neuronal molecular alterations reflecting these behavioral outcomes. Since alterations in neurofilament in sepsis patients have been associated with cognitive dysfunction (Ehler et al., 2017; Ehler et al., 2019), we first determined the levels of non-phosphorylated and hyperphosphorylated neurofilament (NF) in the brain of control and septic mice using IF analysis. We found that post-sepsis cognitive dysfunction was associated with a profound decrease (∼40% reduction) in unphosphorylated NF (Figure 6A, B, SMI-32 immunopositivity) and a robust surge in hyperphosphorylated (p)NF (4.6±0.1fold, Figure 6C, D, SMI-31 immunoreactivity) in both hippocampus and cortex (Figure 6-1 A-D), 1 month after recovery. These alterations in NF have been found to precede neurodegeneration (Wilson et al., 2016) (Krasemann et al., 2017) and cognitive decline (Bussière et al., 2003; Thangavel et al., 2009; Vickers et al., 1994) both in aging and Alzheimer’s disease. These results suggest the presence of axonal injury in our model and identify NF and pNF as promising neuronal molecular markers of the long-term functional repercussions of severe systemic inflammation.

**Figure 6:**
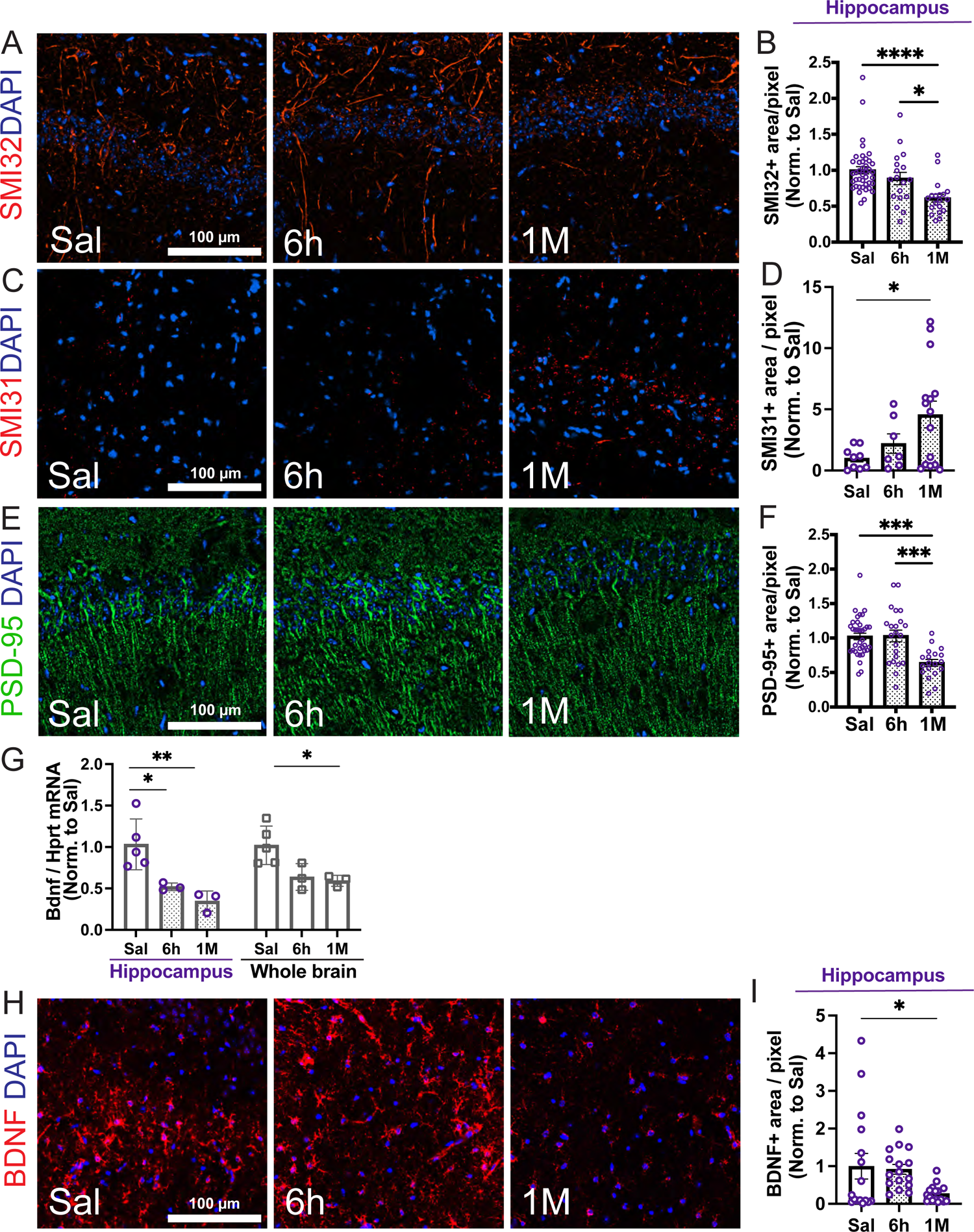
Systemic inflammation induces alterations in neurofilament (NF) phosphorylation, post-synaptic density protein 95 (PSD-95) and brain derived neurotrophic factor (BDNF) in hippocampus. Representative immunofluorescence images and quantification for (A, B) SMI32 (unphosphorylated NF, red channel), (C, D) SMI31 (hyperphosphorylated NF, pNF, red channel) and (E, F) PSD-95 (green channel) staining in the hippocampus from Saline-treated controls, 6 hours (acute phase) and 1 month (chronic phase) post LPS administrations. Systemic inflammation caused a decrease in SMI32 and PSD-95, together with an increase in SMI31 immunopositivity in hippocampus. Representative images from CA1 region are shown. (G) Decrease in Brain derived neurotrophic factor (Bdnf) mRNA levels in both hippocampus and whole brain following systemic inflammation (acute, 6h, and chronic, 1-month (M) phases). RT-qPCR analyses of Bdnf mRNA normalized by Hprt are shown. Fold change LPS compared to saline is plotted. Data points represent the average of duplicates or triplicates of each mouse. Individual values and average ± SEM are shown. N=3-5 mice/group. (H) Representative immunofluorescence images for BDNF staining (red channel) in the hippocampus from Saline-treated controls, 6 hours and 1 month post Saline/LPS administration. (I) Quantitative analysis representing the total area of BDNF+ staining in the hippocampus. (A, C, E, H) Representative pictures (20x magnification). Scale bar=100μm. Nuclear staining (DAPI) is shown in the blue channel. (B, D, F, I) Quantification of the positive SMI32, SMI31, PSD-95 and BDNF areas are shown. Each data point represents one region of interest (ROI) normalized by the mean of Saline. N=3-5 mice/group. The individual values and the mean±SEM are shown. ****p<0.0001, one-way ANOVA followed by Tukey’s test.

Synapse loss and reduction of synapse markers have been reported to be hallmark features of memory impairment (Shao et al., 2011). Thus, we examined the levels of post-synaptic density protein 95 (PSD-95), a major post-synaptic scaffold protein which plays a key role in the structural and functional integrity of excitatory synapses (Chen et al., 2008; El-Husseini Ael et al., 2002). We found that immunoreactivity for PSD-95 was significantly decreased in hippocampus (∼36% reduction, Figure 6E-F) and cortex (∼60% reduction, Figure 6-1 E, F), 1 month after recovery from systemic inflammation. These data suggests that the described systemic inflammation-induced cognitive impairment model leads to synapsis damage and loss.

We also aimed to investigate the alterations in the expression of the neurotrophic factor brain derived neurotrophic factor (BDNF), which play a key role in hippocampal neuronal function (Heldt et al., 2014) (Hofer et al., 1990). mRNA quantification by qRT-PCR analyses showed a ∼40% and 65% reduction in Bdnf mRNA at 1 month following LPS injection (p<0.05) in hippocampus and whole brain, respectively (Figure 6G). These results were further validated by histopathological analyses, showing a sustained depletion in the BDNF positive area in both hippocampus (72% reduction, Figure 6H-I) and cortex at 1 month (80% decrease, Figure 6-1 G, H).

Altogether, these data indicate that our model of systemic inflammation-induced cognitive impairment leads to alterations in neurofilament, PSD-95 and BDNF in the chronic phase (1 month after recovery), suggesting the induction of diffuse axonal damage, post-synapsis degeneration and impaired neurotrophism.

## DISCUSSION

In the current study we have established and standardized a murine model of systemic inflammation-induced cerebral microvascular dysfunction and cognitive impairment. This model consists of three intraperitoneal injections of bacterial endotoxin (LPS), which cause an acute and rapid systemic hyperinflammatory response that resolves approximately at 72 hours after the last administration. This acute phase is characterized by functional (albumin and small-molecule BBB leakage) and molecular alterations (induction of PLVAP and PAI-1) in the cerebral microvasculature and neuroinflammation. Remarkably, some of these perturbations persisted after recovery from the systemic inflammatory response (chronic phase), including small molecule BBB leakage, elevated levels of PAI-1 in cerebral microvessels, peri/intravascular fibrin/fibrinogen deposition and microglial activation. In addition, the chronic phase was also characterized by changes in neuronal molecular markers indicative of axonal and synaptic damage (alterations in neurofilaments and PSD-95), and impaired neurotrophism (decreased BDNF levels), which were associated with memory impairment. This standardized model and the identified molecular markers of neurovascular dysfunction will permit future investigations of the mechanisms underlying systemic inflammation-induced cognitive decline, in particular, elucidation of the role of the cerebral microvasculature and its therapeutic potential.

In our investigation, we used the endotoxemia model to recapitulate key clinical features of systemic inflammatory response syndrome (SIRS) and sepsis, such as the generation of an acute systemic hyper-inflammatory response *via* the activation of pathogen recognition receptors, and the release of pro-inflammatory cytokines (Banks et al., 2015; Ho et al., 2015; Pires et al., 2020; Seemann et al., 2017; Zhao et al., 2019). We monitored the progression of the inflammatory response by quantifying the clinical signs of systemic inflammation (mMSS), body temperature and plasma IL-6 levels, a reliable biomarker of systemic inflammation and sepsis-related morbidity. In our model, this systemic inflammatory response progressed quickly after the first LPS injection, which is in line with previous studies (Seemann et al., 2017) and rapidly resolved 3 days after the last administration. Notably, systemic inflammation induced chronic lasting effects in the brain beyond recovery, which were reflected in the impaired spatial, recognition and emotional memory in LPS-treated mice. The hippocampus, in coordination with the cortex, plays a critical role in learning and memory, and is particularly vulnerable to acute and chronic neuroinflammation. Sepsis survivors suffer from compromised spatial memory, decreased visual attention, impaired executive functions within the first year, and a persistent reduction in hippocampal and cortical volume up to 2 years after hospital discharge (Andonegui et al., 2018; Gunther et al., 2012; Semmler et al., 2013). Similar results have been reported in rodents, showing impaired learning and hippocampal damage after sepsis (Basak et al., 2021; Ge et al., 2023; Lee et al., 2008; Semmler et al., 2008). Based on existing evidence and the observed deficits in spatial, recognition and contextual memory, our results suggest that compromised hippocampal and cortical neuronal function is recapitulated in our model. One limitation of our study is the use of bacterial endotoxin instead of an active pathogen; therefore, the endotoxemia model is rather a model of sterile SIRS. Nevertheless, SIRS, in response to onco-therapies or trauma, has also been associated with cognitive dysfunction(Gust et al., 2017), and the systemic inflammatory response and cognitive deficits after recovery were faithfully recapitulated in our model. Another limitation of our study was that we did not monitor changes in the peripheral leukocyte populations. Instead, our main objective was to focus on the molecular and functional neurovascular alterations and their relationship with the systemic inflammatory response. For this reason, we phenotypically characterized the kinetics of the systemic response based on the clinical signs (MSS system) and the plasma IL-6 levels, which is a reliable well-established biomarker of systemic inflammation and sepsis morbidity(Remick et al., 2002; Shapiro et al., 2010; Zhang et al., 2013). According to our findings, the described scoring system is a noninvasive and effective way of tracking the systemic inflammatory response, showing kinetics similar to that of IL-6.

Our sepsis/SIRS model caused chronic lasting effects on the brain after the resolution of the systemic inflammatory response which were reflected in memory deficits and molecular changes indicative of diffuse axonal injury, synapse damage, impaired neurotrophism and neuroinflammation. Alterations in neurofilament, PSD-95 and brain-derived neurotrophic factor (BDNF) were key molecular changes identified in our study that were associated with cognitive impairment 1 month after recovery from systemic inflammation. Neurofilaments are major components of the axonal cytoskeleton. We observed a marked decrease in the levels of the nonphosphorylated NF heavy subunit (NF-H) and an increase in the levels of hyperphosphorylated (pNF-H), which is specifically transported and accumulated in disconnected axons following injury (Anderson et al., 2008; Kelberman et al., 2022; Shaw et al., 2005). As these components are released into the cerebrospinal fluid (CSF) and later into the systemic circulation (Yuan et al., 2012), and because of their easy access and feasibility for sampling across patients, most of the research concerning pNF-H has focused on circulating levels in the blood and CSF as a predictive biomarker of neuronal injury in different pathological conditions, including Alzheimer’s disease (Boylan et al., 2013; Darras et al., 2019; Gatson et al., 2014; Hu et al., 2002; Lee et al., 2020; Shaw, 2015; Shibahashi et al., 2016; Zurek et al., 2012). However, similar alterations in NF have been described in the brain (Bussière et al., 2003; Thangavel et al., 2009; Vickers et al., 1994) and retina (Wilson et al., 2016); pre-clinical and clinical data indicate that hyperphosphorylation of neurofilaments precedes degeneration (Wilson et al., 2016) and that decreased immunoreactivity for nonphosphorylated NF in hippocampal and cortical neurons is linked to cognitive impairment in aging and dementia in patients (Bussière et al., 2003; Thangavel et al., 2009; Vickers et al., 1994). In addition to axonal injury, synapse loss and reduction in synapse markers are also hallmark features of post-sepsis memory impairment (Huerta et al., 2016) and early dementia (Shao et al., 2011; Terry et al., 1991). Phagocytosis of post-synaptic proteins by activated microglia has been proposed as a cellular mechanism underlying post-synaptic degeneration in dementia (Roy et al., 2020) and neuroinflammatory pathologies (Jafari et al., 2021). In our model, reduction in PSD-95 levels, a major postsynaptic scaffold protein that plays a key role in the structural and functional integrity of excitatory synapses (Chen et al., 2008; El-Husseini Ael et al., 2002) along with microglial activation were associated with sepsis-induced cognitive dysfunction. We also found decreased mRNA and protein levels of BDNF in the brain and hippocampus, both in the acute phase and 1 month after recovery from systemic inflammation. Neurotrophic factors play a crucial role in neuronal function and differentiation. BDNF is a major regulator of neuronal growth and synapse plasticity and is closely involved in learning and memory formation(Chen et al., 2006; Dincheva et al., 2014; Giza et al., 2018; Soliman et al., 2010). This neurotrophic factor is expressed both in cortical and subcortical regions, with the highest expression in hippocampal neurons (Bathina et al., 2015; Hofer et al., 1990; Timmusk et al., 1993). Previous studies have shown that inflammation downregulates BDNF expression, in both rodents and humans (Bouvier et al., 2017; Garbossa et al., 2022; Gibney et al., 2013; Pedard et al., 2021; Yap et al., 2021). At the same time, BDNF limits the neuroinflammatory response in the brain (Parrott et al., 2021), (Wu et al., 2020). Although additional research is needed to explore the signaling pathways leading to the changes observed in BDNF, as well as NF and PSD-95, and to clarify the precise functional implications of these changes in our model, our findings have nonetheless highlighted key molecular neuronal markers that are associated with cognitive dysfunction following severe systemic inflammation.

Another key contribution of our study was the identification of functional and molecular alterations in the cerebral microvasculature in our sepsis model during and after recovery from the systemic inflammatory response. We found robust opening of the BBB to plasma proteins (albumin) and small molecules during the acute phase, which coincided with the upregulation of PLVAP, a caveolar protein that plays a key role in non-specific endothelial transcytosis and vascular leakage (Bosma et al., 2018; Wisniewska-Kruk et al., 2016). In contrast to endothelial cells from peripheral organs, PLVAP expression is actively repressed in the cerebrovascular and retinal endothelium (Daneman et al., 2010). Our data are consistent with recent studies reporting the induction of PLVAP in hypoxic-ischemic and inflammatory pathological conditions and its role in blood-brain and blood-retinal barrier leakage(Bosma et al., 2018; Callegari et al., 2023; Munji et al., 2019; Shue et al., 2008; Wisniewska-Kruk et al., 2016). We postulate that this early opening of the BBB in the acute phase of sepsis allows the leakage of plasma proteins and cytokines (Beurel et al., 2009; Winkler et al., 2017) (Qin et al., 2007), which initiates microglial activation (Tejera et al., 2019), impairing their homeostatic function(Jafari et al., 2021) and contributing to synapse loss (Huerta et al., 2016), a hallmark of cognitive impairment and early dementia (Shao et al., 2011; Terry et al., 1991). Another important finding in the acute phase in our model was the upregulation of the antifibrinolytic PAI-1 in the cerebral microvasculature. Increased levels of PAI-1 in sepsis impairs fibrin degradation and contributes to intravascular fibrin deposition, microthrombus formation and microvascular dysfunction(Shapiro et al., 2010; Tipoe et al., 2018). In the healthy brain and cerebral microvasculature, PAI-1 is expressed at minimal levels (Sutton et al., 1994; Yamamoto et al., 2005), yet it is induced in patients with stroke(Callegari et al., 2023; Tjärnlund-Wolf et al., 2012) and Alzheimer’s disease (Oh et al., 2014) (Angelucci et al., 2023; Melchor et al., 2005). Remarkably, in addition to these acute changes observed in the cerebral microvasculature, our study revealed key alterations that continued after recovery from the systemic inflammatory response. We found persistent small molecule (cadaverine) BBB leakage and elevated levels of both PAI-1 mRNA and protein along with increased intra/perivascular fibrin/fibrinogen deposition in the chronic phase. These novel findings suggest that chronic cerebral microvascular dysfunction after recovery from systemic inflammation, namely, small molecule leakage and intra/perivascular coagulation, could further exacerbate neurovascular inflammation, synaptic and axonal degeneration and the progression of cognitive dysfunction (Griemert et al., 2019; Liu et al., 2011; Zipser et al., 2007) (Merlini et al., 2019; Petersen et al., 2018). In addition, they highlight the therapeutic potential of the endothelium not only in the acute phase but also in the chronic phase after recovery from sepsis. Interestingly, in other pathologies such as Alzheimer’s disease (Angelucci et al., 2023; Eruysal et al., 2023; Oh et al., 2014) and neurodegenerative diseases (Griemert et al., 2019; Liu et al., 2011; Zipser et al., 2007) (Merlini et al., 2019; Petersen et al., 2018), BBB leakage (Nation et al., 2019) and increased levels of PAI-1 and fibrinogen, have also been found to correlate with cognitive dysfunction and disease severity. Our data indicate that our SIRS/sepsis model faithfully recapitulates key features of cerebral microvascular dysfunction (that is, BBB leakage and thromboinflammation) in the acute and chronic phases, which are associated with cognitive impairment, providing a validated model to study the contributions of the cerebral microvasculature to sepsis-associated cognitive decline.

In conclusion, the present study provides a standardized and validated animal model for examining of the impact of systemic inflammation on cerebral microvascular and cognitive dysfunction. The molecular and functional alterations identified in the cerebral microvasculature suggest the transition to a permeability and procoagulant state, which persisted after recovery from the systemic inflammatory response. Our work has uncovered molecular markers of cerebral microvascular dysfunction in sepsis highlighting promising opportunities for the therapeutic targeting of the endothelium in the acute and chronic phases. This experimental model will permit future investigations of the mechanisms underlying systemic inflammation-induced cognitive dysfunction, such as the role of the cerebral microvasculature, its therapeutic potential and identification of novel endothelial therapeutic targets and biomarkers. These future studies will be critical to improve our understanding of the mechanisms by which systemic inflammation affects the development and progression of cognitive impairment and other neurodegenerative conditions, including dementia or Alzheimeŕs disease, so that novel effective therapies for these conditions can be developed.

## ACKNOWLEDGEMENTS

Funding sources: NIH, NINDS 5R01NS114561 to TS, R01MH12315 to FSL, R00MH119320 to HM.

## EXTENDED DATA FIGURE AND TABLE LEGENDS

**Figure 2-1. Alteration of each parameter of Murine Sepsis Score (MSS) following LPS injection** This figure illustrates the changes observed in each individual parameter that contributes to the overall MSS score, subsequent to LPS administration. Parameters such as Appearance, Level of Consciousness, Activity, Response to Stimulus, Respiration, and Nesting behavior are plotted over time to detail the acute (Inj 1 to 72h) and sub-acute (96h to 1 week, wk) effects of LPS-induced systemic inflammation. The mean of individual value±SEM are shown. N=7 Saline, N=13 LPS.

**Figure 3-1. Alterations in microvascular inflammation and integrity post systemic inflammation in cortex and hippocampus.** (A-D) Histopathological validation and quantification of BBB leakage in cortex. N=5-8/group. (A, B) Albumin leakage in cortex using Alexa-594 albumin as a tracer shows an increase in albumin+ area at 6h following LPS administration that returns to baseline levels at 1 month. (C, D) Alexa-555 Cadaverine leakage showed an increase in BBB permeability in cortex 6h after LPS treatment that persists at 1 month. Panels A and C show representative pictures (40x magnification) of the cortex with a 2x digital zoom inset. Scale bar=50μm. (E, F) Histopathological validation and quantification of PAI-1 alterations (red channel) in the cortical microvessels (Glut1 positive areas, green channel). N=3 mice/group. (G-J) Fibrin/ogen (FBG) immunofluorescence analyses and quantification in intra/perivascular areas (Glut1 positive areas, green channel) in the cortex (G, H) and hippocampus (I, J). N=3 mice/group. Panels E, G and I show representative pictures (20x magnification), along with 2x-zoom insets for panels G and I. Scale bar=100μm. Nuclear staining (DAPI) is shown in the blue channel. (B, D, F, H, J) Each data point represents one region of interest (ROI) from the scanned image. Data are fold induction normalized to the mean of Saline. The individual values and the mean±SEM are shown. *p<0.05, **p<0.005, ***p<0.0005, ****p<0.0001, one-way ANOVA followed by Tukey’s test.

**Figure 4-1. Systemic inflammation induces the expression of neuroinflammatory markers in cortex.** Immunofluorescence images and quantification for (A, B) GFAP, (C, D) IBA1, (E, F) P2Y12 and (G, H) P2Y12/IBA1 staining in the parenchyma of cortex from Saline-treated controls, and 6 hours and 1 month post Saline/LPS administration (all nuclei labeled with DAPI, blue channel). Panels A, C, E and G show 20x representative pictures. Scale bar=100μm. (B, D, F, H) Quantitative analysis representing the total area of GFAP+, IBA1+, P2Y12+ and IBA1+P2Y12+ staining in cortex. Notice the increase in GFAP, IBA-1 and IBA1/P2Y12 immunopositivity and the decrease in P2Y12 in the acute phase, suggesting astrocyte and microglial activation. Each data point symbolizes one region of interest (ROI) from the scanned image and was normalized by the mean of Saline. N=3-5 mice/group. The individual values and the mean±SEM are shown. *p<0.05, **p<0.005, ***p<0.0005, ****p<0.0001, one-way ANOVA followed by Tukey’s test.

**Figure 6-1. Systemic inflammation induces alterations in neurofilament phosphorylation, post-synaptic density 95 (PSD-95) and brain derived neurotrophic factor (BDNF) in cortex.** Immunofluorescence images and quantification for (A, B) SMI32 (unphosphorylated NF, red channel), (C, D) SMI31 (hyperphosphorylated NF, pNF, red channel), (E, F) PSD-95 (green channel) and (G, H) BDNF (red channel) in the cortex of Saline-treated controls, and 6 hours and 1 month post Saline/LPS administrations. Nuclear staining (DAPI) is shown in all images (blue channel). Scale bar=100μm. Notice the decrease in SMI32, PSD-95 and BDNF, together with an increase in SMI31 immunopositivity in cortex after LPS injections. Panels A, C, E and G show representative pictures (20x magnification) and B, D, F and H show the quantitative analyses. Each data point symbolizes one region of interest (ROI) from the scanned sections, normalized by the mean of Saline. N=3-5 mice/group. The individual values and the mean±SEM are shown. **p<0.005, ***p<0.0005, ****p<0.0001, one-way ANOVA followed by Tukey’s test.

**Extended data figure supporting** figures 3, 4 and 6**. ROI used for quantification of histological analyses.** Schematic of the ROI used for quantification.

**Extended Data Table 1. Statistical table for all analyses.**

