## Supplemental Figures and Table for "Molecular and functional alterations in the cerebral microvasculature in an optimized mouse model of sepsis-associated cognitive dysfunction"

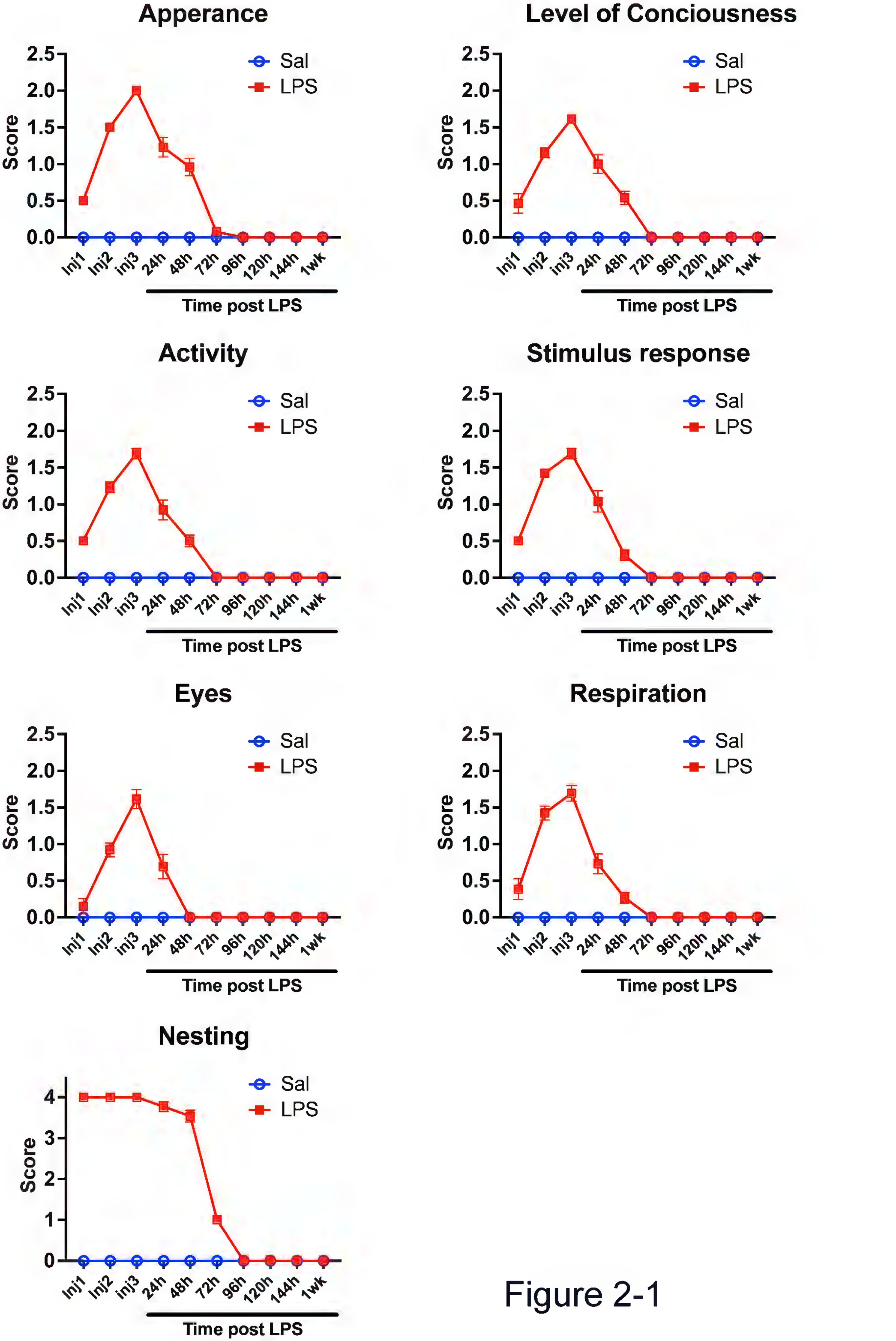

Apperance
Level of Conciousness
Activity
Stimulus response
Eyes
Respiration
Nesting
Score
Sal
LPS
Inj1
Inj2
inj3
24h
48h
72h
96h
120h
144h
1wk
Time post LPS


Figure 2-1

Figure 3-1

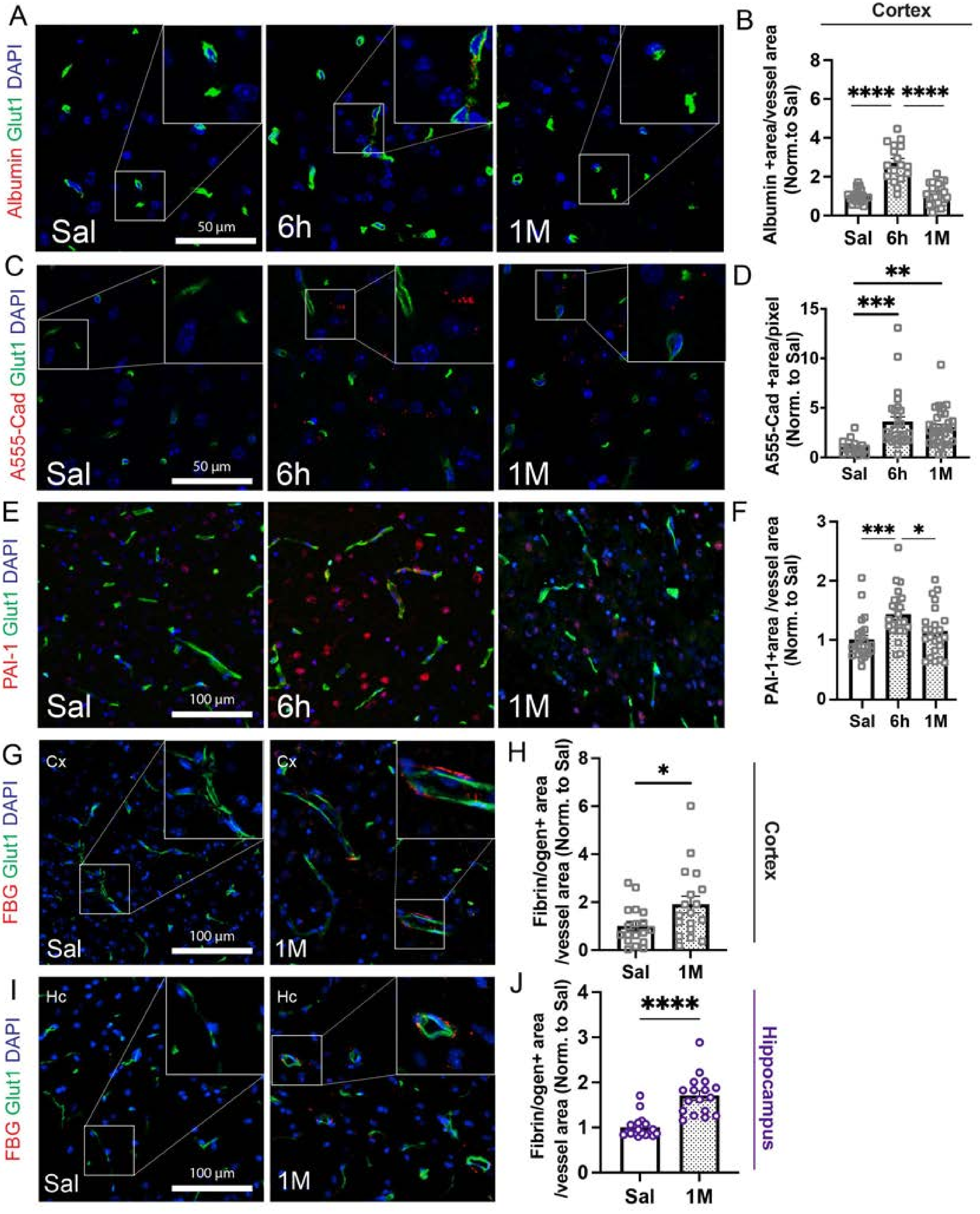

A
Albumin Glut1 DAPI
Sal
50 µm
6h
1M
B
Cortex
Albumin +area/vessel area (Norm. to Sal)
8
6
4
2
0
****
****
Sal
6h
1M
C
A555-Cad Glut1 DAPI
Sal
50 µm
6h
1M
D
A555-Cad +area/pixel (Norm. to Sal)
15
10
5
0
**
***
Sal
6h
1M
E
PAI-1 Glut1 DAPI
Sal
100 µm
6h
1M
F
PAI-1+area /vessel area (Norm. to Sal)
3
2
1
0
***
*
Sal
6h
1M
G
FBG Glut1 DAPI
Cx
Sal
100 µm
Cx
1M
H
Fibrin/ogen+ area /vessel area (Norm. to Sal)
8
6
4
2
0
*
Sal
1M
Cortex
I
FBG Glut1 DAPI
Hc
Sal
100 µm
Hc
1M
J
Fibrin/ogen+ area /vessel area (Norm. to Sal)
4
3
2
1
0
****
Sal
1M
Hippocampus

### Figure 4-1

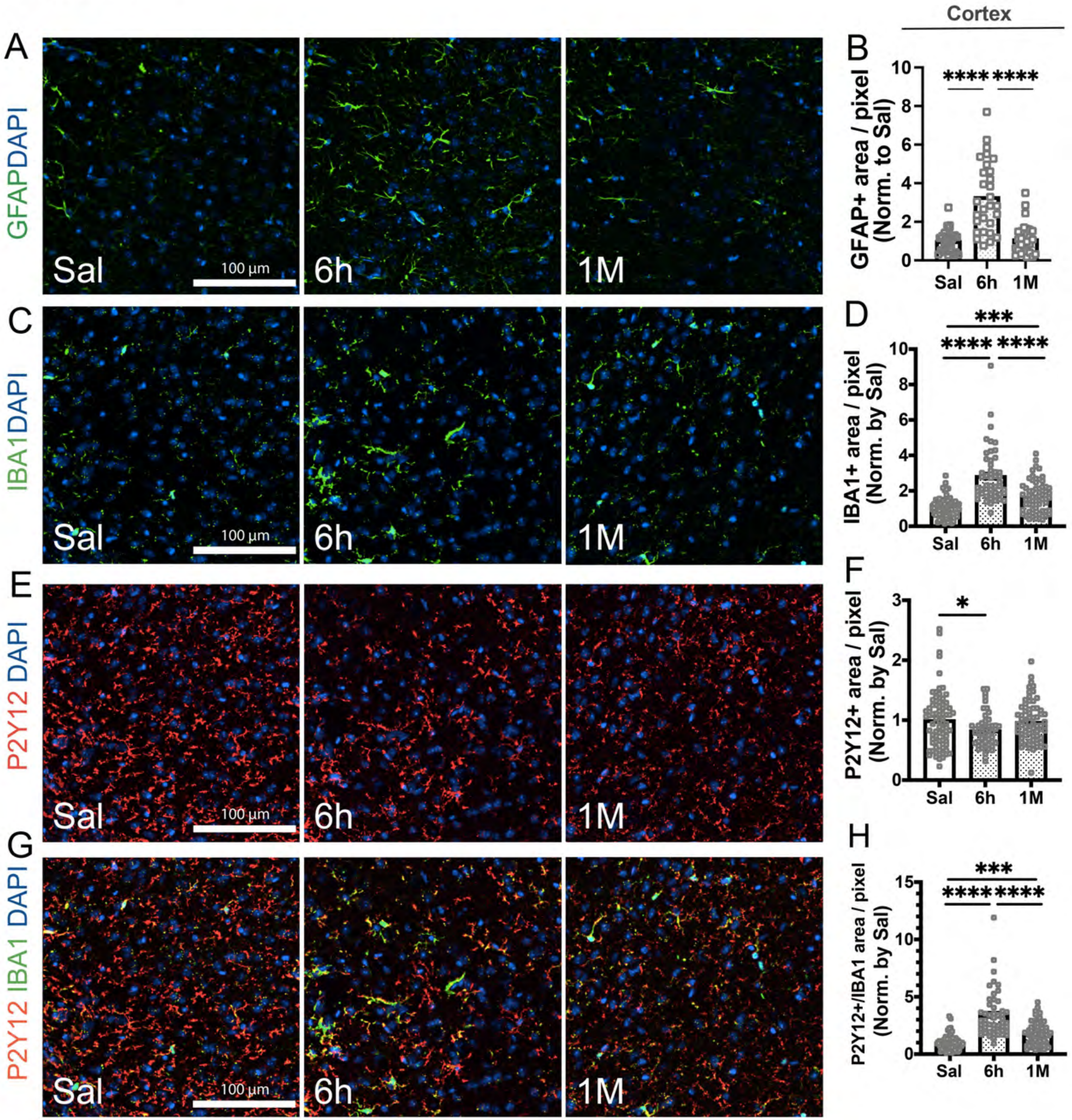

Cortex
A
GFAPDAPI
Sal
100 µm
6h
1M
B
GFAP+ area / pixel (Norm. to Sal)
****
****
Sal
6h
1M
C
IBA1DAPI
Sal
100 µm
6h
1M
D
IBA1+ area / pixel (Norm. by Sal)
***
****
****
E
P2Y12 DAPI
Sal
100 µm
6h
1M
F
P2Y12+ area / pixel (Norm. by Sal)
*
G
P2Y12 IBA1 DAPI
Sal
100 µm
6h
1M
H
P2Y12+/IBA1 area / pixel (Norm. by Sal)
***
****
****

Figure 6-1

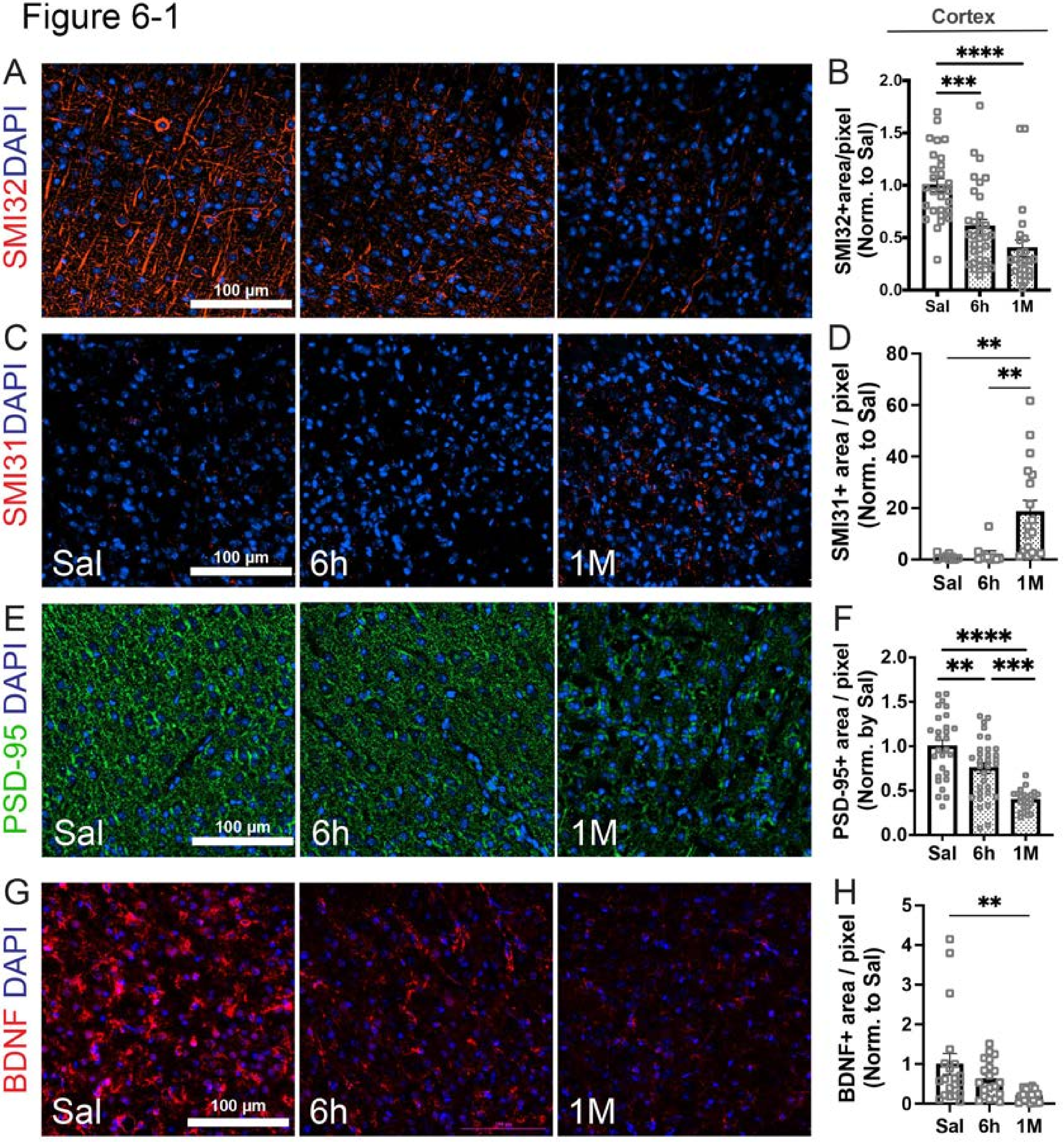

Cortex
A
SMI32DAPI
100 µm
B
SMI32+area/pixel (Norm. to Sal)
Sal
6h
1M
C
SMI31DAPI
D
SMI31+ area / pixel (Norm. to Sal)
E
PSD-95 DAPI
F
PSD-95+ area / pixel (Norm. by Sal)
G
BDNF DAPI
H
BDNF+ area / pixel (Norm. to Sal)

Figure 6-2
(Extended Data Supporting Figures 3, 4, 6)

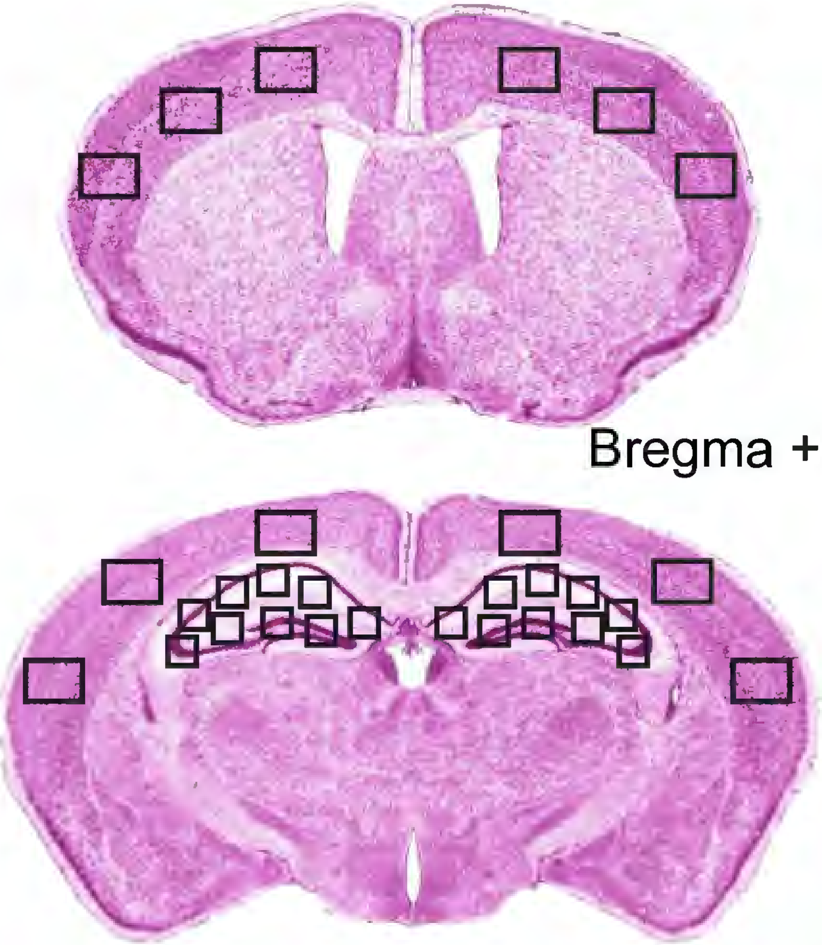

Bregma +0.5 to -0.3 mm

Bregma -1.8 to -2.6 mm

### Statistical Table

| Figure | Data Structure | Sample Size | Test Used | Degree of Freedom & p-value |
|---|---|---|---|---|
| Fig. 2 (modified Murine Sepsis Score) | Normal Distribution | 200 | Two-way ANOVA | 190, p<0.0001 |
| Fig. 2 (Body Temperature) | Normal Distribution | 200 | Two-way ANOVA | 190, p<0.0001 |
| Fig. 2 (IL-6 plasma levels) | Normal Distribution | 26 | One-way ANOVA | 25, p<0.0001 |
| Fig. 2 (Body Weight) | Normal Distribution | 200 | Two-way ANOVA | 190, p=0.002 |
| Fig. 3 (BBB Leakage) | Normal Distribution | 20 | One-way ANOVA | 19, p<0.0001 |
| Fig. 3 (Alexa- 594 Albumin Hippocampus) | Normal Distribution | 56 | One-way ANOVA | 55, p<0.0001 |
| Fig. 3 (Alexa-555 Cadaverine Hippocampus) | Normal Distribution | 66 | One-way ANOVA | 65, p<0.0001 |
| Fig. 3 (Cldn5 qPCR MV) | Normal Distribution | 15 | One-way ANOVA | 14, p=0.0009 |
| Fig. 3 (Plvap qPCR MV) | Normal Distribution | 15 | One-way ANOVA | 14, p<0.0001 |
| Fig. 3 (Cav1 qPCR MV) | Normal Distribution | 15 | One-way ANOVA | 14, p=0.0727 |
| Fig. 3 (Serpine1 qPCR MV) | Normal Distribution | 15 | One-way ANOVA | 14, p=0.0077 |
| Fig. 3 (PAI1 Hippocampus) | Normal Distribution | 120 | One-way ANOVA | 119, p<0.0001 |
| Fig. 4 (GFAP Hippocampus) | Normal Distribution | 120 | One-way ANOVA | 119, p<0.0001 |
| Fig. 4 (IBA1 Hippocampus) | Normal Distribution | 119 | One-way ANOVA | 118, p=0.0009 |
| Fig. 4 (P2Y12 Hippocampus) | Normal Distribution | 119 | One-way ANOVA | 118, p=0.0180 |
| Fig. 4 (P2Y12/IBA1 Hippocampus) | Normal Distribution | 119 | One-way ANOVA | 118, p=0.0002 |
| Fig. 5 (Distance Traveled) | Normal Distribution | 76 | Two-way ANOVA | 72, p=0.2906 |
| Fig. 5 (Primary Latency) | Normal Distribution | 76 | Two-way ANOVA | 72, p=0.0361 |
| Fig. 5 (Primary Latency Day4) | Normal Distribution | 19 | Unpaired t-test | 17, p=0.0132 |
| Fig. 5 (Total Exploration Time) | Normal Distribution | 26 | Unpaired t-test | 24, p=0.2290 |
| Fig. 5 (Discrimination Ratio) | Normal Distribution | 26 | Unpaired t-test | 24, p=0.0165 |
| Fig. 5 (Baseline Motion) | Normal Distribution | 41 | Unpaired t-test | 39, p=0.7968 |
| Fig. 5 (Baseline Motion) | Normal Distribution | 205 | Two-way ANOVA | 200, p=0.5687 |
| Fig. 5 (Contextual Fear Recall) | Normal Distribution | 41 | Unpaired t-test | 39, p=0.0206 |
| Fig. 6 (SMI32 Hippocampus) | Normal Distribution | 78 | One-way ANOVA | 77, p=0.0002 |
| Fig. 6 (SMI31 Hippocampus) | Normal Distribution | 31 | One-way ANOVA | 30, p=0.0342 |
| Fig. 6 (PSD95 Hippocampus) | Normal Distribution | 65 | One-way ANOVA | 64, p=0.0006 |
| Fig. 6 (Bdnf Hippocampus qPCR) | Normal Distribution | 11 | One-way ANOVA | 10, p=0.0210 |
| Fig. 6 (Bdnf Whole brain qPCR) | Normal Distribution | 11 | One-way ANOVA | 10, p=0.0067 |
| Fig, 6 (BDNF Hippocampus immunofluorescence) | Normal Distribution | 48 | One-way ANOVA | 47, p=0.0373 |
| Fig. 3-1 (Alexa- 594 Albumin Cortex) | Normal Distribution | 61 | One-way ANOVA | 40, p<0.0001 |
| Fig. 3-1 (Alexa-555 Cadaverine Cortex) | Normal Distribution | 74 | One-way ANOVA | 73, p=0.001 |
| Fig. 3-1 (PAI1 Cortex) | Normal Distribution | 72 | One-way ANOVA | 71, p=0.0011 |
| Fig. 3-1 (Fibrin/ogen Cortex) | Normal Distribution | 37 | Unpaired t-test | 35, p=0.0499 |
| Fig. 3-1 (Fibrin/ogen Hippocampus) | Normal Distribution | 35 | Unpaired t-test | 33, p<0.0001 |
| Fig. 4-1 (GFAP Cortex) | Normal Distribution | 87 | One-way ANOVA | 86, p<0.0001 |
| Fig. 4-1 (IBA1 Cortex) | Normal Distribution | 198 | One-way ANOVA | 197, p<0.0001 |
| Fig. 4-1 (P2Y12 Cortex) | Normal Distribution | 198 | One-way ANOVA | 197, p=0.0376 |
| Fig. 4-1 (P2Y12/IBA1 Cortex) | Normal Distribution | 198 | One-way ANOVA | 197, p<0.0001 |
| Fig. 6-1 (SMI32 Cortex) | Normal Distribution | 84 | One-way ANOVA | 83, p<0.0001 |
| Fig. 6-1 (SMI31 Cortex) | Normal Distribution | 37 | One-way ANOVA | 36, p=0.0019 |
| Fig. 6-1 (PSD95 Cortex) | Normal Distribution | 84 | One-way ANOVA | 83, p<0.0001 |
| Fig. 6-1 (BDNF Cortex) | Normal Distribution | 60 | One-way ANOVA | 59, p=0.0054 |
